# The ventral hippocampus and nucleus accumbens underlie long-term social memory about female conspecifics in male mice

**DOI:** 10.1101/2024.01.08.574751

**Authors:** Akiyuki Watarai, Kiyoshi Ishida, Teruhiro Okuyama

## Abstract

For many social animals, including humans, the ability to remember and recognize conspecifics (i.e., having social memory), especially a mating partner, is essential for adaptive reproductive strategies. Our previous studies have shown that social memory about same-sex conspecifics, namely male-to-male social memory, is stored in hippocampal ventral CA1 (vCA1) neurons and that neural projections from the vCA1 to the nucleus accumbens (NAc) are crucial for social discriminatory behavior. However, how the memory of an opposite-sex conspecific, specifically male-to-female social memory, and how the social memory modulates sexual behavior in males remain unknown. Herein, we investigated the ultrasonic vocalizations (USVs) and socially approaching behavior of males toward a female before and after social memory formation. We found that the total duration of USVs was reduced when the male mice interacted with a familiar female. This reduction in USV emission was blocked by optogenetic inhibition of vCA1 neurons, as well as D1- and D2-dopamine receptor (D1R and D2R) expressing neurons in the NAc. Furthermore, fiber photometry recordings in the NAc revealed that the activities of both D1R and D2R neuronal populations were altered during social interaction with familiar or novel females after the process of familiarization. The response of D1R-neurons was reduced only when interacting with familiar females. In contrast, although responses of D2R-neurons were reduced toward both familiar and novel females after social memory formation, the reduction towards novel females was significantly more pronounced compared to that towards familiar females. Our findings suggested that the activities of both vCA1 neurons and D1R- and D2R-expressing NAc neurons in males were differentially modulated by the presence of social memory about females during adaptive reproductive strategies.

**Significance Statement:** Long-term social memory about females modulates male sexual behavior, specifically ultrasonic vocalization emissions, mediated by hippocampal ventral CA1 and dopamine receptor expressing neurons in the nucleus accumbens.

## Introduction

Mammals exhibit species-specific reproductive strategies adapted to their respective environments. Monogamous male prairie voles prefer to socially interact with a mating partner, but polygamous male mice do not show such preference for mating females (Beery et al., 2018). Furthermore, in polygamous rodents, repeated exposure to the same female mating partner results in the attenuation of sexual behaviors, and motivation for the sexual behaviors is restored upon encountering an unfamiliar female (Fiorino et al., 1997). Regardless of the reproductive strategy, male rodents have the ability to distinguish a familiar female from a novel one and modify their sexual behaviors accordingly (Wilson et al., 1963; Walum and Young, 2018). Therefore, the modification of sexual behaviors can be utilized to assess the degree of the formation of social memory about females in male rodents. Upon encountering a female, male rodents exhibit a sequential progression of sexual behaviors from investigative sniffing to mounting, ultimately leading to intromission and ejaculation. Throughout all these behavioral phases until the ejaculation, males emit ultrasonic vocalizations (USVs) (Sales, 1972; Nyby, 1983). While investigative sniffing and mounting have also been observed in male-male interactions, sexually mature males emit USVs only in the presence of females (Karigo et al., 2021). Additionally, the emission of USVs is frequently and robustly observed in male-female interactions and sexual behavior compared to other steps of sexual behaviors such as mounting, intromission, and ejaculation, and most of the emitted USVs originate from males (Wang et al., 2008). Thus, as it is easy to record and determine the source of USVs, we attempted to quantify USV emissions as a useful parameter for assessing the social memory of male mice about female conspecifics.

We previously demonstrated that the hippocampal ventral CA1 (vCA1) subregion is crucial for storing social memories about same-sex conspecific individuals, with no involvement in non-social object memory (Okuyama et al., 2016). Male mice reduce investigative sniffing toward a male conspecific as they form and consolidate their social memory about the individual, and the modulation of social discriminatory behavior is mediated by the excitatory projection from the vCA1 to the nucleus accumbens (NAc) (Okuyama et al., 2016). Given that male and female information is primarily processed in overlapping brain regions, albeit with potential cellular-level differences (Li and Dulac, 2018), we hypothesized that the functions of the vCA1 and NAc are necessary to modulate sexual behavior based on the formation and consolidation of social memory about females. The NAc is involved in the modulation of behavior based on memory and learning, with contributions not only from the predominant D1- and D2-dopamine receptor (D1R and D2R) expressing neurons, but also from calcium/calmodulin-dependent protein kinase II (CaMKII) (Zhou et al., 2019; Jackson et al., 2016; Amaral et al., 2021). In prairie voles, the essential role of the NAc is implicated in pair bond (Edwards and Self, 2006; Aragona and Wang, 2009). D2R signaling is necessary for pair bonding with a mating partner, and D1R expression is upregulated afterward (Edwards and Self, 2006; Aragona and Wang, 2009), which is suggested to contribute to the behavioral change after pair bonding (Loth and Donaldson, 2021). Furthermore, it has been reported that mating leads to an increase in the number of neurons in the NAc that preferentially respond to the mating partner; however, the specific cell type of these neural populations has not been defined (Scribner et al., 2020). Therefore, it remains unclear which neural populations in the male NAc function to modulate sexual behavior based on social memory about females.

In this study, we recorded USV emissions from male mice to evaluate their social memory about females in response to a familiar and novel female. After isolating the males, we exposed them to each female and found that they emitted fewer USVs toward the familiar female compared to the novel one. This specific reduction of USVs to the familiar female was impaired by optogenetic inhibition of the vCA1 during the reunion with the familiarized female. Furthermore, when we optogenetically inhibited D1R- and D2R-expressing NAc neurons, but not CaMKII, male mice showed impaired specific reduction of USVs to the familiar female. By measuring the calcium transient of D1R- and D2R-expressing NAc neurons before and after familiarization using fiber photometry, we observed differential activation of these neuronal populations in response to the familiar and novel female. These results suggest that the neural activities of D1R- and D2R-expressing NAc neurons in males are differentially modulated by the presence of social memory about females, contributing to the elicitation of adaptive sexual behaviors.

## Result

### Males emit fewer USVs in the presence of a familiar female than a novel oneC

We first attempted to establish a novel behavioral paradigm in male mice to quantify the degree of social memory about opposite-sex female conspecifics by quantifying the emission of USVs. For assessing long-term social memory about females, we recorded the USVs emitted by the test-male toward a C3H/HeJ (C3H) female and a C57BL/6J (B6) female as well as social approaching behavior of the test-male toward these females before and after the 2 hours of familiarization with the C3H female (***Figure 1A, 1B***). During familiarization, the C3H female was restrained by a pencil holder to prevent the initiation of mating behavior as USV emission is significantly reduced after mating. After the recording, the generic features of USVs were detected by an automated algorithm (USVSEG, Tachibana et al., 2020). The total duration of USVs emitted toward the familiarized female (C3H), but not the unfamiliarized novel female (B6), was significantly reduced after the familiarization (***Figure 1C***: before vs after, Tukey–Kramer post hoc comparisons: B6, p = 0.72: C3H, p < 0.01). In contrast, the total duration of the social approaching behavior seemed not to be affected by the familiarization (***Figure 1D***: before vs after, Tukey–Kramer post hoc comparisons: B6, p = 0.09: C3H, p = 0.41).

**Figure 1.**
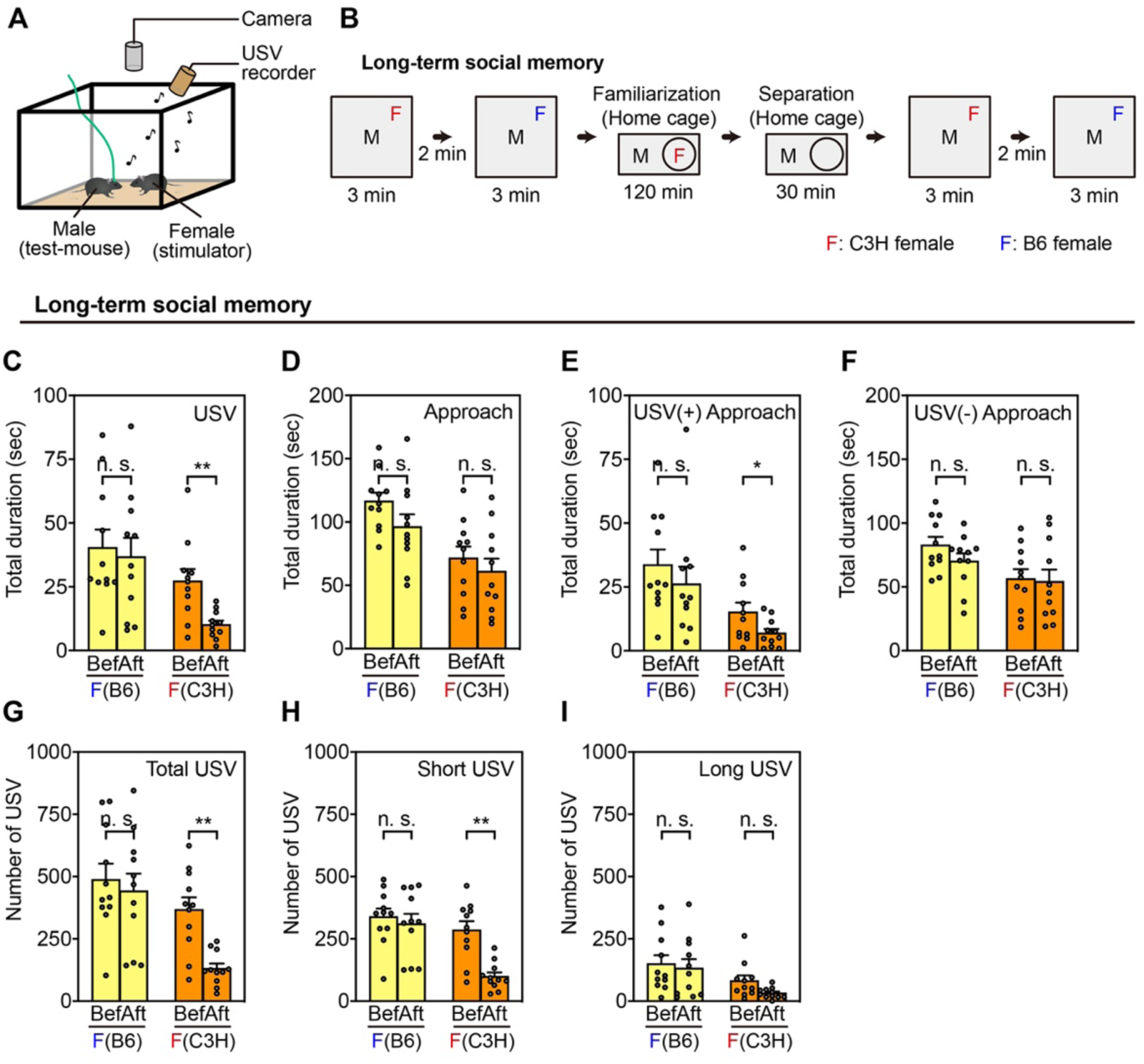
Modulation of Sexual Behaviors in the Long-term Social Memory Task. **(A)** Setup of ultrasonic vocalization (USV) recordings for quantifying social memory. (**B)** Schematic view of the behavioral paradigm. USVs and approaching behaviors by a male mouse (black M) toward C3H (red F) and B6 (blue F) stimulator female mice were recorded before and after 2-hour familiarization with the C3H female in the home cage. During the familiarization phase, the C3H female was confined inside the pencil holder to prevent mating. **(C, D)** Total duration of USVs **(C)** and approaching behaviors **(D)** by the male toward B6 (left) and C3H (right) females. **p<0.01. **(E, F)** Total duration of approaching behaviors with **(E)** and without **(F)** the presence of USVs by the male toward B6 (left) and C3H (right) females. *p<0.05. **(G–I)** The number of total **(G)**, short **(H)**, and long **(I)** USVs emitted by the male toward B6 (left) and C3H (right) females. **p<0.01. **(C–I)** n = 11 mice. Two-way repeated-measure analysis of variance (rANOVA) followed by Tukey–Kramer post hoc comparison.

A recent study showed that female- and male-directed mounting behaviors are distinguishable by the presence or absence of USVs (Karigo et al., 2021). Thus, we distinguished the two types of social approaching behaviors by the presence or absence of USVs and found that only the approaching behavior accompanied with USVs was modulated by social memory (before vs after, Tukey–Kramer post hoc comparisons; ***Figure 1E***, USV (+) approach: B6, p = 0.32: C3H, p < 0.05; ***Figure 1F***, USV (-) approach: B6, p = 0.20: C3H, p = 0.83). Furthermore, we analyzed the components of the recorded USVs, since USVs are composed of a complex mixture of long USVs (above 50 msec) and short USVs (below 50 msec). The number of emitted total USVs toward the familiar female was reduced after the familiarization (***Figure 1G***: before vs after, Tukey–Kramer post hoc comparisons: B6, p = 0.63: C3H, p < 0.01), corresponding with the total duration of USVs shown in ***Figure 1C***, which was mainly ascribed to the reduced number of short USVs (before vs after, Tukey–Kramer post hoc comparisons; ***Figure 1H***, short USVs: B6, p = 0.61: C3H, p < 0.01; ***Figure 1I***, long USVs: B6, p = 0.71: C3H, p = 0.06).

### Optogenetic inhibition of vCA1 neurons impairs the reduction of emitted USVs toward a familiar female

Next, to examine whether vCA1 neurons are essential for long-term social memory about opposite-sex female conspecifics, we performed the long-term social memory task with optogenetic inhibition of vCA1 neurons. To specifically target vCA1 pyramidal neurons, the adeno-associated virus (AAV), AAV5-EF1α:DIO-eArch3.0-EYFP, or its control virus, AAV5-EF1α:DIO-EYFP, was bilaterally injected into the vCA1 of Trpc4-Cre males (Okuyama et al., 2016), and the vCA1 was also the target of the optic fiber implants (***Figure 2A***). We confirmed that the expression of eArch-EYFP was abundant in the vCA1 (***Figure 2B–2D***). Optogenetic inhibition of vCA1 neurons was performed during the social interaction after the familiarization (i.e., recall phase) (***Figure 2E***) and the USVs and approaching behavior were analyzed (***Figure 2 and Figure 2–figure supplement 1***) similar to the previous experiments with untreated mice (***Figure 1***). Although the emitted USVs toward the familiar C3H female, but not toward the novel B6 female, were significantly reduced both in the control and vCA1 inhibition groups (***Figure 2F***: before vs after, Tukey–Kramer post hoc comparisons: EYFP-B6, p = 0.58: EYFP-C3H, p < 0.01: eArch-B6, p = 0.27: eArch-C3H, p < 0.01), the levels of reduction measured by the discrimination index were significantly impaired by optogenetic inhibition of vCA1 neurons (***Figure 2J***: EYFP vs eArch, Tukey–Kramer post hoc comparisons: B6, p = 0.42: C3H, p < 0.05). In contrast, the emitted USVs toward the novel B6 female were not reduced in either the control or the vCA1 inhibition groups after the familiarization with the C3H female (***Figure 2F***), corresponding with unaffected discrimination indices (***Figure 2J***).

**Figure 2.**
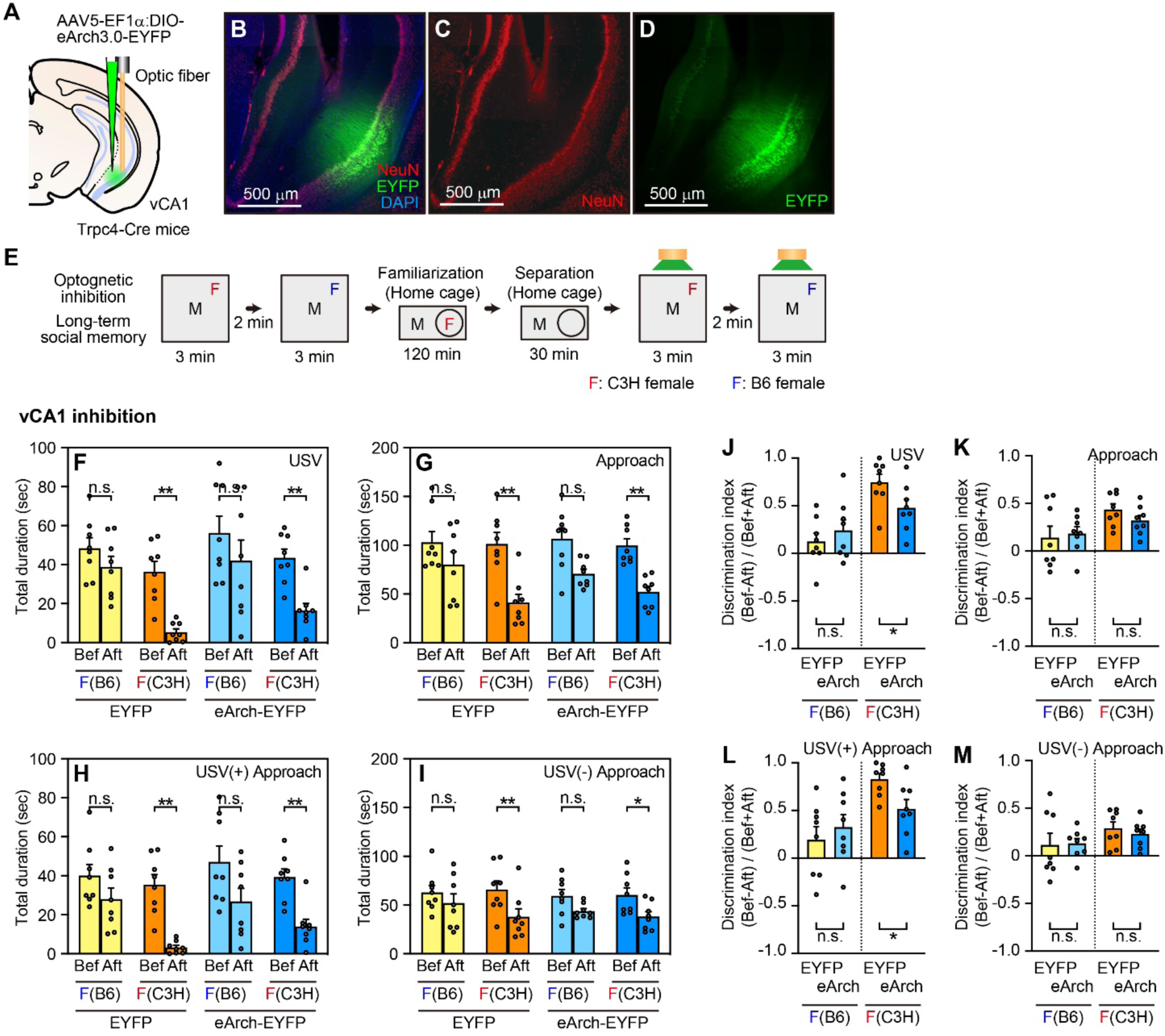
Optogenetic Inhibition of the vCA1 During the Long-term Memory Task. **(A)** Viral injection (AAV5-EF1α:DIO-eArch3.0-EYFP) and optic fiber implantation into the vCA1 of Trpc4-Cre mice. **(B–D)** Representative images of vCA1 expressing DAPI (blue), NeuN (red), and EYFP (green). **(E)** Schematic view of the behavioral paradigm with optogenetic inhibition. The inhibition was conducted during the recording after familiarization. **(F, G)** Total duration of USVs (**F**) and approaching behaviors (**G**) by the male with vCA1 inhibition (left, control group, EYFP; right, inhibition group, eArch-EYFP) toward the B6 and C3H females (left and right side of each group, respectively). *p<0.05; **p<0.01. **(H, I)** Total duration of approaching behaviors with (**H**) and without (**I**) USVs by the male with vCA1 inhibition (left, control group, EYFP; right, inhibition group, eArch-EYFP) toward the B6 and C3H females (left and right side of each group, respectively). *p<0.05; **p<0.01. **(J–M)** Comparison of the discrimination index corresponding to **(F–I)**. *p<0.05. **(F-M)** EYFP, n = 8; eArch-EYFP, n = 8 mice. **(F–I)** Before vs After familiarization, Three-way rANOVA followed by Tukey–Kramer post hoc comparison. **(J–M)** EYFP vs, eArch-EYFP, Two-way rANOVA followed by Tukey–Kramer post hoc comparison.

In addition, we analyzed the total duration of approaching behavior (***Figure 2G***: before vs after, Tukey– Kramer post hoc comparisons: EYFP-B6, p = 0.60: EYFP-C3H, p < 0.01: eArch-B6, p = 0.24: eArch-C3H, p < 0.01; ***Figure 2K***: EYFP vs eArch, Tukey–Kramer post hoc comparisons: B6, p = 0.77: C3H, p = 0.17), and the duration of approaching behavior with and without the presence of USVs (USV (+) approach: ***Figure 2H***: before vs after, Tukey– Kramer post hoc comparisons: EYFP-B6, p = 0.59: EYFP-C3H, p < 0.01: eArch-B6, p = 0.18: eArch-C3H, p < 0.01; ***Figure 2L***: EYFP vs eArch, Tukey–Kramer post hoc comparisons: B6, p = 0.50: C3H, p < 0.05; USV (-) approach: ***Figure 2I***: before vs after, Tukey–Kramer post hoc comparisons: EYFP-B6, p = 0.76: EYFP-C3H, p < 0.01: eArch-B6, p = 0.52: eArch-C3H, p < 0.05; ***Figure 2M***: EYFP vs eArch, Tukey–Kramer post hoc comparisons: B6, p = 0.91: C3H, p = 0.48). Unlike the previous experiments using untreated wild-type (WT) males (***Figure 1***), the familiarization with the C3H female reduced both the total duration of approaching behavior and the duration of approaching behavior without the presence of USVs. However, the comparison between control and inhibition groups based on the discrimination index revealed that vCA1 inhibition did not have a significant effect on the duration of these behaviors (***Figure 2K and 2M***). Notably, the reduction in approaching behavior accompanied by a USVs as a result of the familiarization was impaired by vCA1 inhibition (***Figure 2L***), further evidencing that the approaching behavior with USVs rely on social memory stored in vCA1.

### D1R- and D2R-expressing NAc neurons are essential for the social memory-dependent reduction of USVs

We previously showed the pivotal function of the vCA1 afferents to the NAc shell for the formation of long-term social memories about same-sex conspecifics (Okuyama et al., 2016). Thus, we attempted to investigate the role of the NAc in social memory concerning opposite-sex conspecifics and identify the cell type of essential NAc medium spiny neurons (MSNs). The NAc contributes to multiple behavioral inhibition and compulsions and is mainly composed of D1R- or D2R-expressing neurons. Thus, to silence D1R- or D2R-expressing neurons, AAV5-EF1α:DIO-eArch3.0-EYFP or AAV5-EF1α:DIO-EYFP was bilaterally injected into the NAc shell of D1R-Cre or D2R-Cre males, and optic fiber implants also targeted the NAc shell (***Figure 3A***). The expression of eArch-EYFP was abundant in the NAc shell, although it was also observed to a lesser extent in the NAc core (***Figure 3B–3I***).

**Figure 3.**
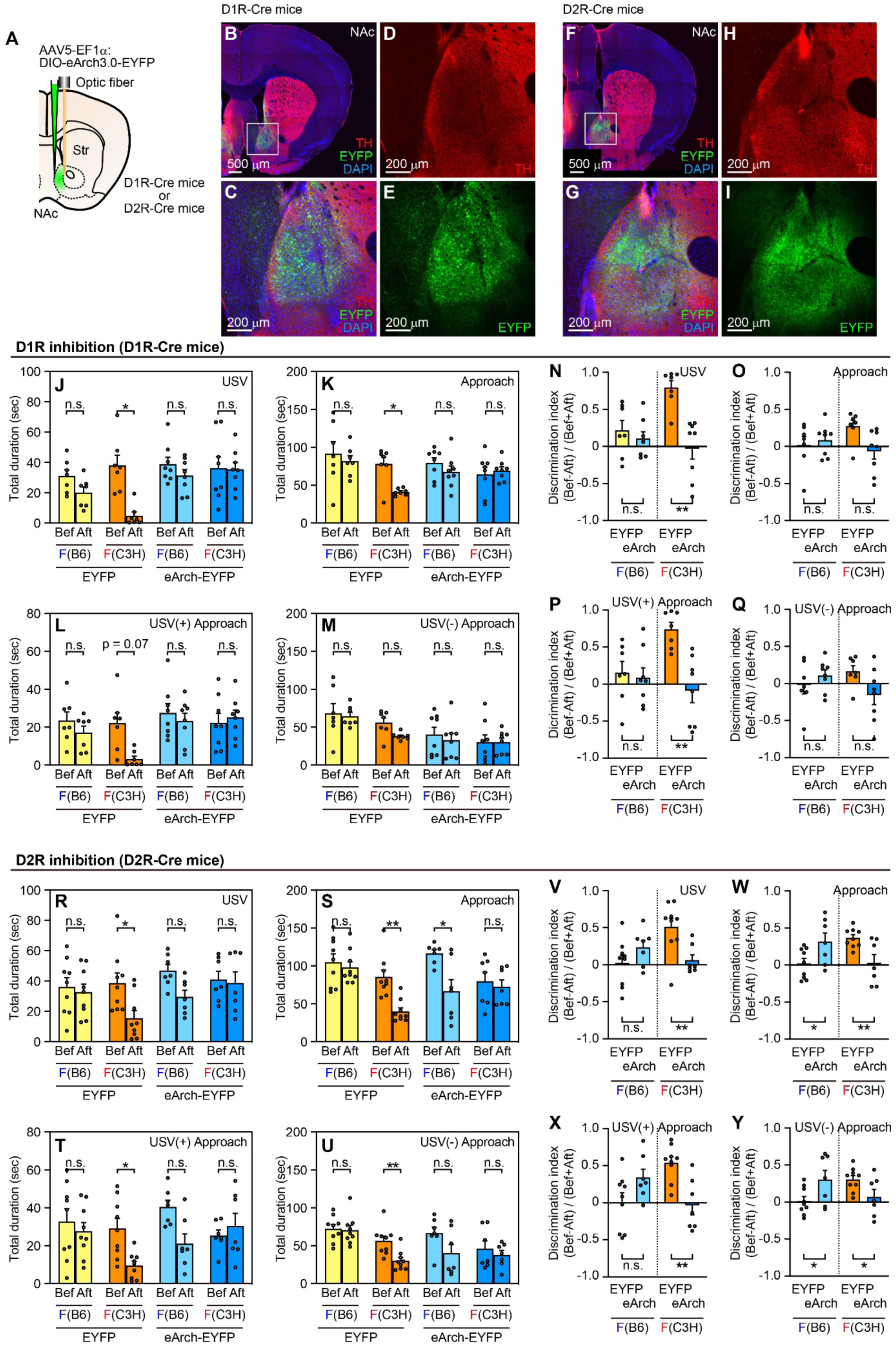
Optogenetic Inhibition of NAc D1R- or D2R-Expressing Neurons During The Long-term Memory Task. **(A)** Viral injection (AAV5-EF1α:DIO-eArch3.0-EYFP) and optic fiber implantation in the NAc of D1R-Cre or D2R-Cre mice. **(B–E)** Representative images of the NAc of D1R-Cre mice expressing DAPI (blue), TH (red), and EYFP (green). Zoomed views of white squares in **(B)** are shown in **(C–E)**. **(F–I)** Representative images of NAc of Drd2-Cre mice expressing DAPI (blue), TH (red), and EYFP (green). Zoomed views of white square in **(F)** are shown in **(G–I)**. **(J, K)** Total duration of USVs **(J)** and approaching behaviors **(K)** by the male with NAc-D1R inhibition (left, control group, EYFP; right, inhibition group, eArch-EYFP) toward the B6 and C3H females (left and right side of each group, respectively). *p<0.05. **(L, M)** Total duration of approaching behaviors with **(L)** and without **(M)** USVs by the male with NAc-D1R inhibition (left, control group, EYFP; right, inhibition group, eArch-EYFP) toward the B6 and C3H females (left and right side of each group, respectively). **(N–Q)** Comparison of the discrimination index corresponding to **(J–M)**. **p<0.01. **(R, S)** Total duration of USVs **(R)** and approaching behaviors **(S)** by the male with NAc-D2R inhibition (left, control group, EYFP; right, inhibition group, eArch-EYFP) toward the B6 and C3H females (left and right side of each group, respectively). *p<0.05; **p<0.01. **(T, U)** Total duration of approaching behaviors with **(T)** and without **(U)** USVs by the male with NAc-D2R inhibition (left, control group, EYFP; right, inhibition group, eArch-EYFP) toward the B6 and C3H females (left and right side of each group, respectively). *p<0.05; **p<0.01. **(V–Y)** Comparison of the discrimination index corresponding to **(R–U)**. *p<0.05; **p<0.01. **(J– Q)** EYFP, n = 7; eArch-EYFP, n = 8 mice. **(R–Y)** EYFP, n = 9; eArch-EYFP, n = 7 mice. **(J–M, R–U)** Before vs After familiarization, Three-way r ANOVA followed by Tukey–Kramer post hoc comparison. **(N–Q, V–Y)** EYFP vs eArch-EYFP, Two-way rANOVA followed by Tukey–Kramer post hoc comparison.

Using the same behavioral experiment paradigm as in the previous vCA1 inhibition experiments, targeted neuronal inhibition was performed during the recall phase. The analysis focused on USVs and social approaching behavior (***Figure 3 and Figure 3–figure supplement 1)***. The reduction in the total duration of USVs towards familiar females as a result of social memory was significantly impaired by the inhibition of D1R-neurons (***Figure 3J***: before vs after, Tukey–Kramer post hoc comparisons: EYFP-B6, p = 0.45: EYFP-C3H, p < 0.05: eArch-B6, p = 0.67: eArch-C3H, p = 1.00), corresponding with the impaired discrimination index in the inhibition group (***Figure 3N***: EYFP vs eArch, Tukey–Kramer post hoc comparisons: B6, p = 0.50: C3H, p < 0.01). In the control group, the effect of familiarization on the duration of approaching behavior with the presence of USVs (***Figure 3L***: before vs after, Tukey–Kramer post hoc comparisons: EYFP-B6, p = 0.79: EYFP-C3H, p = 0.07: eArch-B6, p = 0.91: eArch-C3H, p = 0.97), as well as the total duration of approaching behavior (***Figure 3K***: before vs after, Tukey–Kramer post hoc comparisons: EYFP-B6, p = 0.90: EYFP-C3H, p < 0.05: eArch-B6, p = 0.82: eArch-C3H, p = 0.97), was not consistent with the previous experiments with untreated WT males (***Figure 1***). However, by comparing the discrimination indices between the control and inhibition groups, only the approaching behavior with the presence of USVs was found to be impaired by the inhibition of D1R-neurons (EYFP vs eArch, Tukey–Kramer post hoc comparisons; ***Figure 3O***, Approach: B6, p = 0.63: C3H, p = 0.13; ***Figure 3P***, USV (+) Approach: B6, p = 0.74: C3H, p < 0.01). In contrast, the duration of approaching behavior without the presence of USVs was not influenced by the inhibition of D1R-neurons (***Figure 3M***: before vs after, Tukey–Kramer post hoc comparisons: EYFP-B6, p = 0.99: EYFP-C3H, p = 0.15: eArch-B6, p = 0.91: eArch-C3H, p = 1.00; ***Figure 3Q***: EYFP vs eArch, Tukey–Kramer post hoc comparisons, B6 p = 0.38, C3H p = 0.06).

Regarding the total duration of USVs, inhibiting D2R-neurons yielded results similar to those obtained from inhibiting D1R-neurons. The reduction in the total USV duration toward a familiar C3H female was significantly impaired in the inhibition group (***Figure 3R***: before vs after, Tukey–Kramer post hoc comparisons: EYFP-B6, p = 0.95: EYFP-C3H, p < 0.01: eArch-B6, p = 0.11: eArch-C3H, p = 0.97; ***Figure 3V***: EYFP vs eArch, Tukey–Kramer post hoc comparisons: B6, p = 0.17: C3H, p < 0.01). On the other hand, inhibiting D2R-neurons during the recall phase exhibited distinct modulation of approaching behavior. The total duration of approaching behavior and the duration of approaching behavior with the presence or absence of USVs were significantly reduced in the control group, and the reduction was significantly impaired by inhibiting D2R-neurons (***Figure 3S–3U and 3W–3Y***, described in more detail in the source data file). Additionally, both the total duration of approaching behavior and the duration of approaching behavior without the presence of USVs toward novel B6 females were significantly reduced in the inhibition group after the familiarization (before vs after, Tukey–Kramer post hoc comparisons; ***Figure 3S***: EYFP-B6, p = 0.96: EYFP-C3H, p < 0.01: eArch-B6, p < 0.05: eArch-C3H, p = 0.94; ***Figure 3U***: EYFP-B6, p = 1.00: EYFP-C3H, p < 0.01: eArch-B6, p = 0.15: eArch-C3H, p = 0.69), corresponding with the enhanced discrimination index (EYFP vs eArch, Tukey–Kramer post hoc comparisons; ***Figure 3W***: B6, p < 0.05: C3H, p < 0.01; ***Figure 3Y***: B6, p < 0.05: C3H, p < 0.05).

### CaMKII-expressing NAc neurons are not necessary for social memory dependent reductions in USV emission

Since CaMKII-expressing NAc neurons are reported to be involved in memory-related tasks (Amaral et al., 2021), we conducted optogenetic inhibition of these neurons during the long-term social memory task described above. We selectively targeted CaMKII-expressing neurons by bilaterally injecting AAV5-CaMKIIa:eArch3.0-EYFP or AAV5-CaMKIIa:EYFP into the NAc shell of WT males, followed by optical fiber implantation into the same region (***Figure 4A–4D***). The target neurons were inhibited during the recall phase of the experimental paradigm, and the analysis of the USVs and approaching behavior was subsequently performed (***Figure 4 and Figure 4–figure supplement 1)***. The total duration of emitted USVs (***Figure 4E***: before vs after, Tukey–Kramer post hoc comparisons: EYFP-B6, p = 0.37: EYFP-C3H, p < 0.05: eArch-B6, p = 0.68: eArch-C3H, p < 0.05), approaching behavior (***Figure 4F***: before vs after, Tukey–Kramer post hoc comparisons: EYFP-B6, p = 0.50: EYFP-C3H, p < 0.05: eArch-B6, p = 0.96: eArch-C3H, p < 0.01), and approaching behavior with the presence of USVs toward the familiar C3H female, but not toward the novel B6 female, were significantly reduced regardless of the inhibition (***Figure 4G***: before vs after, Tukey–Kramer post hoc comparisons: EYFP-B6, p = 0.75: EYFP-C3H, p <0.05: eArch-B6, p = 0.93: eArch-C3H, p < 0.05). Moreover, there was no significant difference in the extent of the reduction between the control and the inhibition group (***Figure 4I–K***, described in more detail in the source data file). The optogenetic inhibition also had no effect on the duration of approaching behavior without the presence of USVs (***Figure 4H***: before vs after, Tukey–Kramer post hoc comparisons: EYFP-B6, p = 0.49: EYFP-C3H, p = 0.79: eArch-B6, p = 0.63: eArch-C3H, p = 0.09; ***Figure 4L***: EYFP vs eArch, Tukey–Kramer post hoc comparisons: B6, p = 0.08: C3H, p = 0.52).

**Figure 4.**
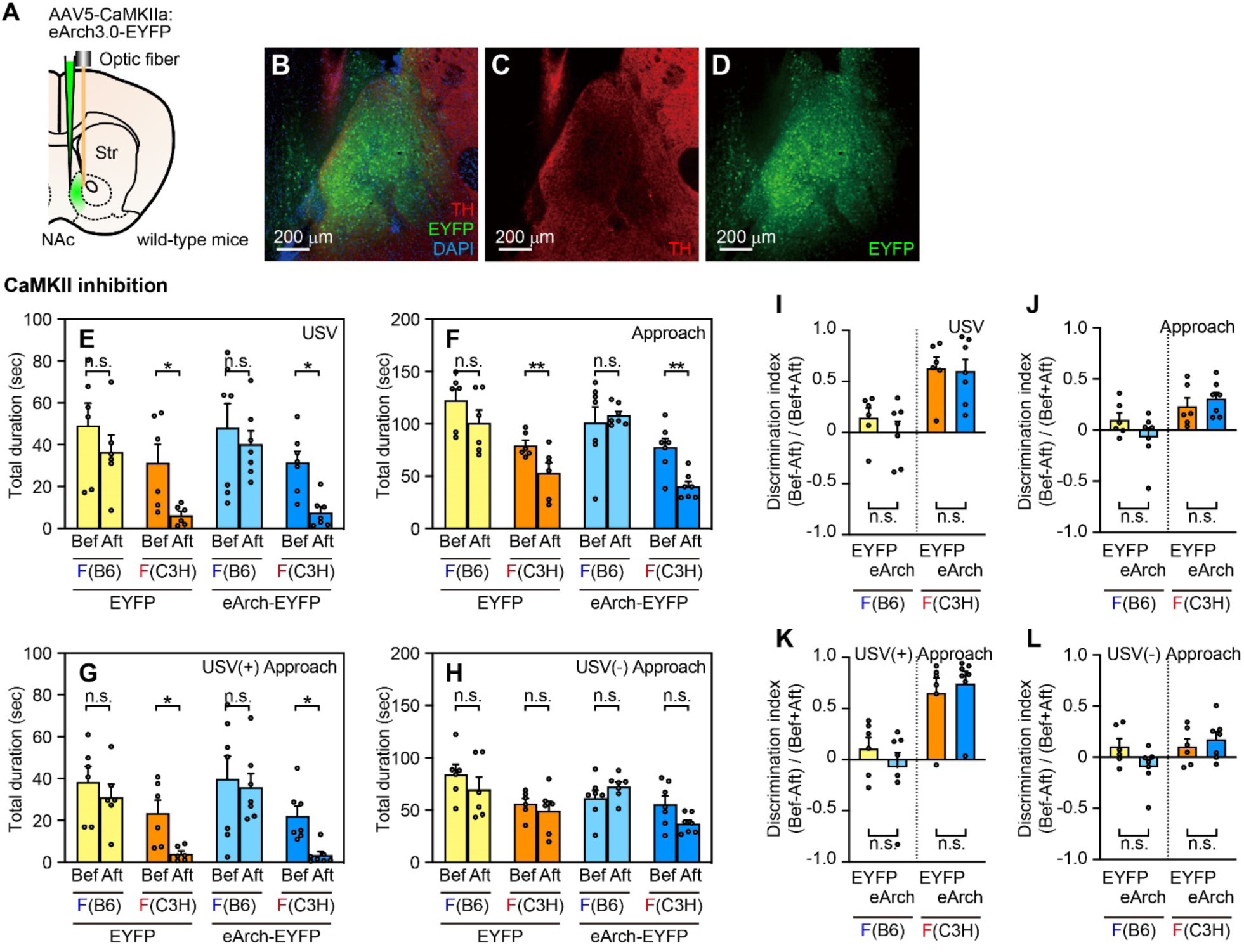
Optogenetic Inhibition of NAc CaMKII-Expressing Neurons During The Long-term Memory Task. Viral injection (AAV5-CaMKIIa:eArch3.0-EYFP) and optic fiber implantation into the NAc of wild type mice. **(B– D)** Representative images of the NAc expressing DAPI (blue), TH (red), and EYFP (green). **(E, F)** Total duration of USVs **I** and approaching behaviors **(F)** by the male with NAc-CaMKII inhibition (left, control group, EYFP; right, inhibition group, eArch-EYFP) toward the B6 and C3H females (left and right side of each group, respectively). *p<0.05; **p<0.01. **(G, H)** Total duration of approaching behaviors with **(G)** and without **(H)** USVs by the male with NAc-CaMKII inhibition (left, control group, EYFP; right, inhibition group, eArch-EYFP) toward the B6 and C3H females (left and right side of each group, respectively). *p<0.05. **(I-L)** Comparison of the discrimination index corresponding to **(E–H)**. **(E–L)** EYFP, n = 6; eArch-EYFP, n = 7 mice. **(E–H)** Before vs After familiarization, Three-way rANOVA followed by Tukey–Kramer post hoc comparison. **(I–L)** EYFP vs eArch-EYFP, Two-way rANOVA followed by Tukey–Kramer post hoc comparison.

Next, we explored the overlaps in the cell types in which we investigated in this study using AAV- and transgenic mice-based approaches. A viral cocktail of Cre-dependent AAV5-hSyn:DIO-histon 2B (H2B)-mCherry, for labeling the D1R- or D2R-expressing neurons, and AAV5-CaMKIIa:EYFP, for visualizing CaMKII-expressing neurons, was injected into the NAc of D1R-Cre and D2R-Cre mice, respectively (***Figure 5A***). In both the injected D1R-Cre (***Figure 5B–5E***) and D2R-Cre (***Figure 5F–5I***) mice, prominent mCherry and EYFP expression were observed. Among the mCherry labeled neurons, 22.6 ± 0.01 (mean ± SEM) percent of D1R-Cre and 20.4 ± 0.01 (mean ± SEM) percent of D2R-Cre mice also expressed EYFP, and there was no significant difference in the ratio of co-labeled neurons in D1R-Cre and D2R-Cre mice (***Figure 5J***: D1R vs D2R, Unpaired t-test, p = 0.24).

**Figure 5.**
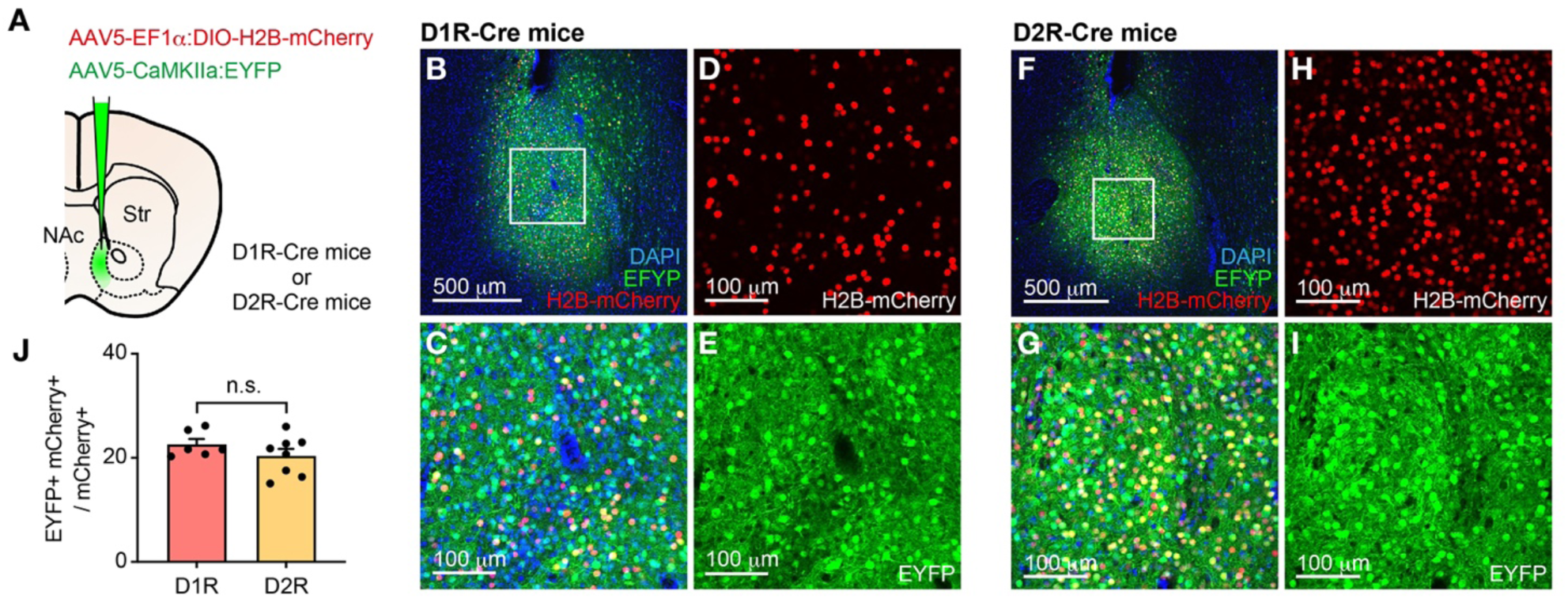
Histological Overlappings of D1R/D2R- and CaMKII-Expressing Neurons. Schematic strategy of EYFP and H2B-mCherry co-expression in the NAc of D1R-Cre or D2R-Cre mice. EYFP and H2B-mCherry are expected to be expressed in CaMKII-expressing and Cre-expressing neurons in the NAc, respectively. **(B–E)** Immunostained confocal images of the NAc of D1R-Cre mice expressing DAPI (blue), EYFP (green), and H2B-mCherry (red). Zoomed views of white squares in **(B)** are shown in **(C–E)**. **(F–I)** Immunostained confocal images of the NAc of D2R-Cre mice expressing DAPI (blue), EYFP (green), and H2B-mCherry (red). Zoomed views of white square in **(F)** are shown in **(G–I)**. **(J)** Ratio of EYFP-mCherry co-expressing neurons to mCherry-expressing neurons in D1R-Cre and D2R-Cre mice. D1R-Cre, n = 6; D2R-Cre, n = 6 samples. p = 0.24, Unpaired t-test.

### *In vivo* physiological activities of D1R- and D2R-expressing neurons in the NAc

To investigate real-time populational activities of D1R-, D2R-, and CaMKII-expressing neurons in the NAc before and after social memory formation, we used fiber photometry technology. The Cre-dependent AAV carrying the fluorescent calcium indicator GCaMP8s (AAV5-syn:FLEX-jGCaMP8s) was stereotaxically injected into the left NAc of D1R-Cre and D2R-Cre male mice, followed by implantation of a 400 μm optical fiber with its tip into the NAc (***Figure 6A***). Additionally, to detect the Ca^2+^ transient of CaMKII-neurons, the mixture of AAV5-syn:FLEX-jGCaMP8s and AAV5-CaMKII:Cre was stereotaxically injected in the left NAc of WT males, followed by the same implantation of an optical fiber (***Figure 6A***). After surgical recovery, we measured the neuronal population activities in the NAc of males during social interaction with female mice by monitoring jGCaMP8s signals in consecutive two days recordings (***Figure 6B***). The 30-second recording sessions, in which the test male mouse and a stimulator C3H or B6 female mouse were placed together and allowed to have direct social interaction in the recording arena, were repeated three times with 4-minute separation periods between each session. These consecutive recording sessions were performed before and after the two hours of familiarization with the C3H female to form the social memory (i.e., Pre- and Post-familiarization recording).

**Figure 6.**
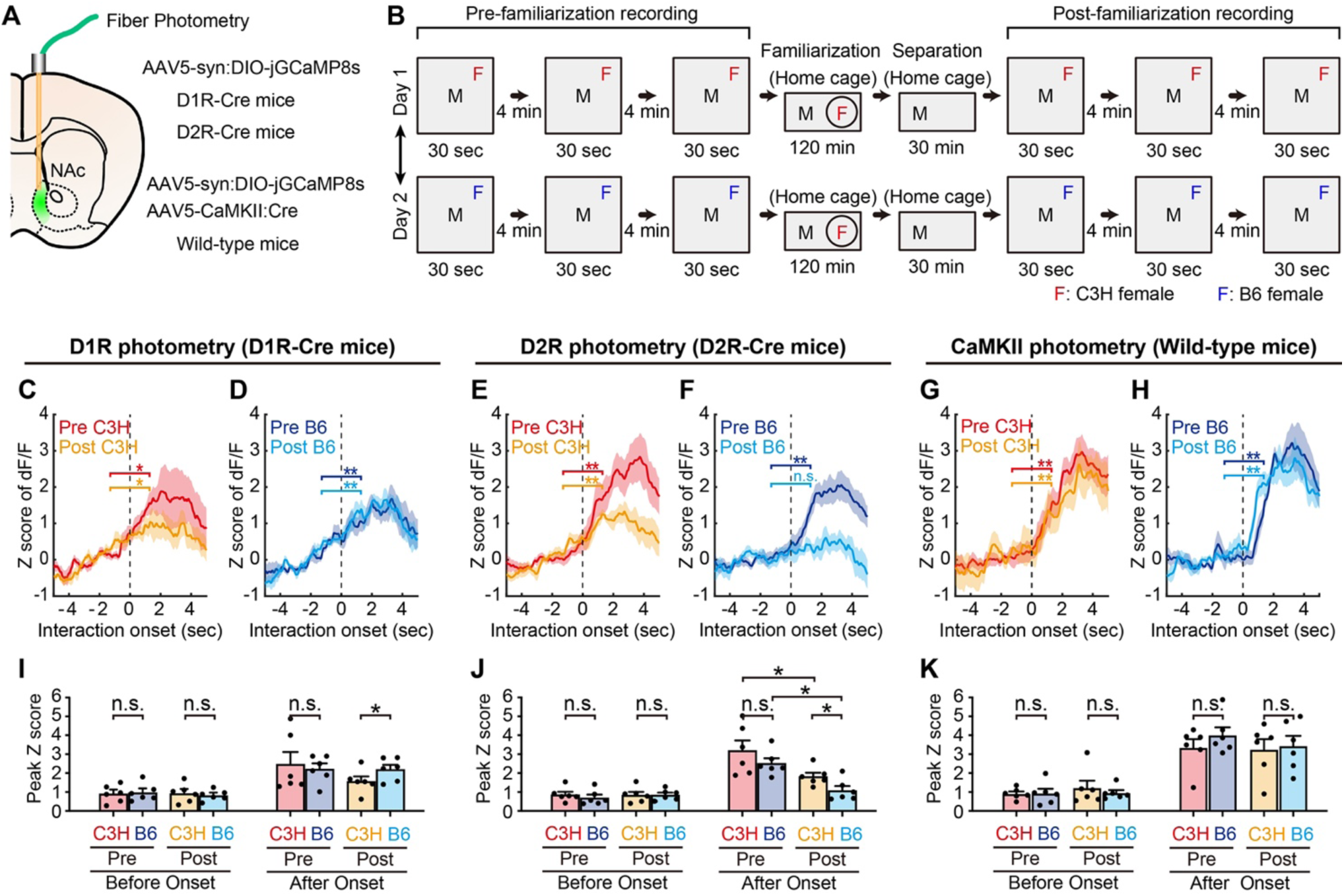
Pre- and Post-Familiarization Calcium Imaging Recordings. **(A)** Schematic strategy of GCaMP expression and optic fiber implantation targeting D1R-, D2R-, and CaMKII-expressing neurons in the NAc. **(B)** Schematic view of the photometry recording paradigm. The recording was conducted over two days. On each day, Ca^2+^ transients during the interaction with either C3H (red F) or B6 (blue F) were recorded before and after the familiarization with a C3H female. **(C, D)** Comparison of Ca^2+^ signals during pre and post familiarization recordings in D1R neuronal population. Ca^2+^ signals recorded five-seconds before and after the interaction onset with C3H **(C)** and B6 **(D)** are shown. *p<0.05; **p<0.01. **(E, F)** Comparison of Ca^2+^ signals during pre and post familiarization recordings in D2R neuronal population. Ca^2+^ signals five-second before and after the interaction onset with C3H **(E)** and B6 **(F)** are shown. n = 6 mice. **p<0.01. **(G, H)** Comparison of Ca^2+^ signals during pre and post familiarization recordings in CaMKII neuronal population. Ca^2+^ signals five-second before and after the interaction onset with C3H **(G)** and B6 **(H)** are shown. *n* = 6 mice. **p<0.01. **I-K** Comparison of the peak amplitude of Ca^2+^ signals before and after the interaction onset, pre and post familiarization, C3H and B6 stimulator corresponding to **(C-H)**. *p<0.05. **(C, D, I)** n = 6 mice. **(E, F, J)** n = 6 mice. **(G, H, K)** n = 6 mice. **(C–K)** Three-way rANOVA followed by Tukey–Kramer post hoc comparison.

It was previously shown that the activity in the ventral tegmental area (VTA) is increased during bouts of social interaction with a novel mouse, with the activation of VTA-NAc projections favoring social interaction. The VTA signals were correlated with onset time of interaction bouts and exhibited habituation over recording epochs (Gunaydin et al., 2014). Thus, to closely examine the activities of the NAc depending on the presence of social memory and eliminate the effect of habituation during the recording, we measured the Ca^2+^ transients within the initial interaction bout during the three sessions of the 30-seconds recording and calculated the average of the three sessions (***Figure 6C–6H***). In the pre-familiarization recording, the area under the curve (AUC) of jGCaMP8s fluorescence observed in D1R-, D2R-, and CaMKII-expressing neurons remarkably increased at the onset of social interaction with both C3H and B6 females (***Figure 6C–6H***: Before Onset vs After Onset, Tukey–Kramer post hoc comparisons: D1R-Pre C3H, p < 0.05: D1R-Pre B6, p < 0.01; D2R-Pre C3H, p < 0.01: D2R-Pre B6, p < 0.01; CaMKII-Pre C3H, p < 0.01, CaMKII-Pre B6, p < 0.01). However, Ca^2+^ transients in the post-familiarization recording showed that the AUC of D2R-expressing neurons was increased only during social interaction with the familiar C3H female (***Figure 6E***: Before Onset vs After Onset, Tukey–Kramer post hoc comparisons: D2R-Post C3H, p < 0.01), but not with the novel B6 female (***Figure 6F***: Before Onset vs After Onset, Tukey–Kramer post hoc comparisons: D2R-Post B6, p = 0.21). The AUC of D1R- or CaMKII-expressing neurons were equivalent both before and after the interaction onset, and during social interaction with C3H and B6 females (***Figure 6C, D, G, and H***: Before Onset vs After Onset, Tukey–Kramer post hoc comparisons: D1R-Post C3H, p < 0.05: D1R-Post B6, p < 0.01; CaMKII-Post C3H, p < 0.01: CaMKII-Post B6, p < 0.01).

Furthermore, to quantitatively compare the interaction between males and C3H and B6 females, we also performed peak analysis (***Figure 6I–6K***). Before the interaction onset, there were no significant differences in the peak Z scored dF/F between familiar and unfamiliar females in either neuronal population (***Figure 6I–6K***: C3H vs B6, Tukey–Kramer post hoc comparisons: D1R-Before Onset Pre, p = 0.677: D1R-Before Onset Post, p = 0.62; D2R-Before Onset Pre, p = 0.54: D2R-Before Onset Post, p = 0.98; CaMKII-Before Onset Pre, p = 0.94: CaMKII-Before Onset Post, p = 0.58). In contrast, the peak Z scored dF/F after the interaction onset in D1R-expressing neurons during the post-familiarization recording was higher in novel B6 females compared to the familiar C3H females (***Figure 6I***: C3H vs B6, Tukey–Kramer post hoc comparisons: D1R-After Onset Pre, p = 0.63: D1R-After Onset Post, p < 0.05), and conversely, it was higher in the familiar C3H females than in B6 females in the case of D2R-expressing neurons (***Figure 6J***: C3H vs B6, Tukey–Kramer post hoc comparisons: D2R-After Onset Pre, p = 0.40: D2R-After Onset Post, p < 0.05). Furthermore, the peak Z scores after the onset in D2R-expressing neurons decreased from the pre-familiarization recording to the post-familiarization recording when interacting with both C3H and B6 females (***Figure 6J***: Pre vs Post, Tukey–Kramer post hoc comparisons: D1R-After Onset B6, p < 0.05: D1R-After Onset C3H, p < 0.05). The peak Z scores after the onset for CaMKII-expressing neurons showed no differences between the pre- and post-familiarization recordings or between the familiar C3H and B6 females (***Figure 6K***: C3H vs B6, Tukey–Kramer post hoc comparisons: CaMKII-After Onset Pre, p = 0.29: CaMKII-After Onset Post, p = 0.82).

### Short-term social memory does not require activity of the vCA1 and NAc

Next, we examined the short-term social memory about opposite-sex conspecifics by assessing the amount of USVs. Direct social interaction with a C3H female for three minutes was repeated five times with two minute inter-trial intervals, followed by the interaction with a novel B6 female for three minutes (***Figure 7A***). Similar to the long-term social memory task (***Figure 1***), we recorded the USVs and approaching behavior (***Figure 7B–7H***). The total USV duration (***Figure 7B***, Tukey–Kramer post hoc comparisons: Trial 1 vs Trial 2, p < 0.05, Trial 1 vs Trial 3, p < 0.01, Trial 1 vs Trial 4, p < 0.01, Trial 1 vs Trial 5, p < 0.01; Trial 5 vs Trial 6, Paired t-test, p < 0.01) and total duration of approaching behavior with the presence of USVs were decayed depending on the presence of social memory (***Figure 7D***, Tukey–Kramer post hoc comparisons: Trial 1 vs Trial 2, p < 0.05, Trial 1 vs Trial 3, p < 0.01, Trial 1 vs Trial 4, p < 0.01, Trial 1 vs Trial 5, p < 0.01; Trial 5 vs Trial 6, Paired t-test, p < 0.01), whereas the total duration of approaching behavior (***Figure 7C***, Tukey–Kramer post hoc comparisons: Trial 1 vs Trial 2, p = 0.34, Trial 1 vs Trial 3, p = 0.06, Trial 1 vs Trial 4, p = 0.25, Trial 1 vs Trial 5, p = 0.31; Trial 5 vs Trial 6, Paired t-test, p < 0.01) and total duration of approaching behavior without USVs were not (***Figure 7E***, Tukey–Kramer post hoc comparisons: Trial 1 vs Trial 2 p = 0.80, Trial 1 vs Trial 3 p = 0.31, Trial 1 vs Trial 4 p = 0.88, Trial 1 vs Trial 5 p = 0.89; Trial 5 vs Trial 6, Paired t-test, p < 0.01). The total number of USVs (***Figure 7F***, Tukey–Kramer post hoc comparisons: Trial 1 vs Trial 2, p < 0.01, Trial 1 vs Trial 3, p < 0.01, Trial 1 vs Trial 4, p < 0.01, Trial 1 vs Trial 5, p < 0.01; Trial 5 vs Trial 6, Paired t-test, p < 0.01), especially the number of short USVs, was also decayed (***Figure 7G***, Tukey–Kramer post hoc comparisons: Trial 1 vs Trial 2 p < 0.01, Trial 1 vs Trial 3 p < 0.01, Trial 1 vs Trial 4 p < 0.01, Trial 1 vs Trial 5 p < 0.01; Trial 5 vs Trial 6, Paired t-test, p < 0.01).

**Figure 7.**
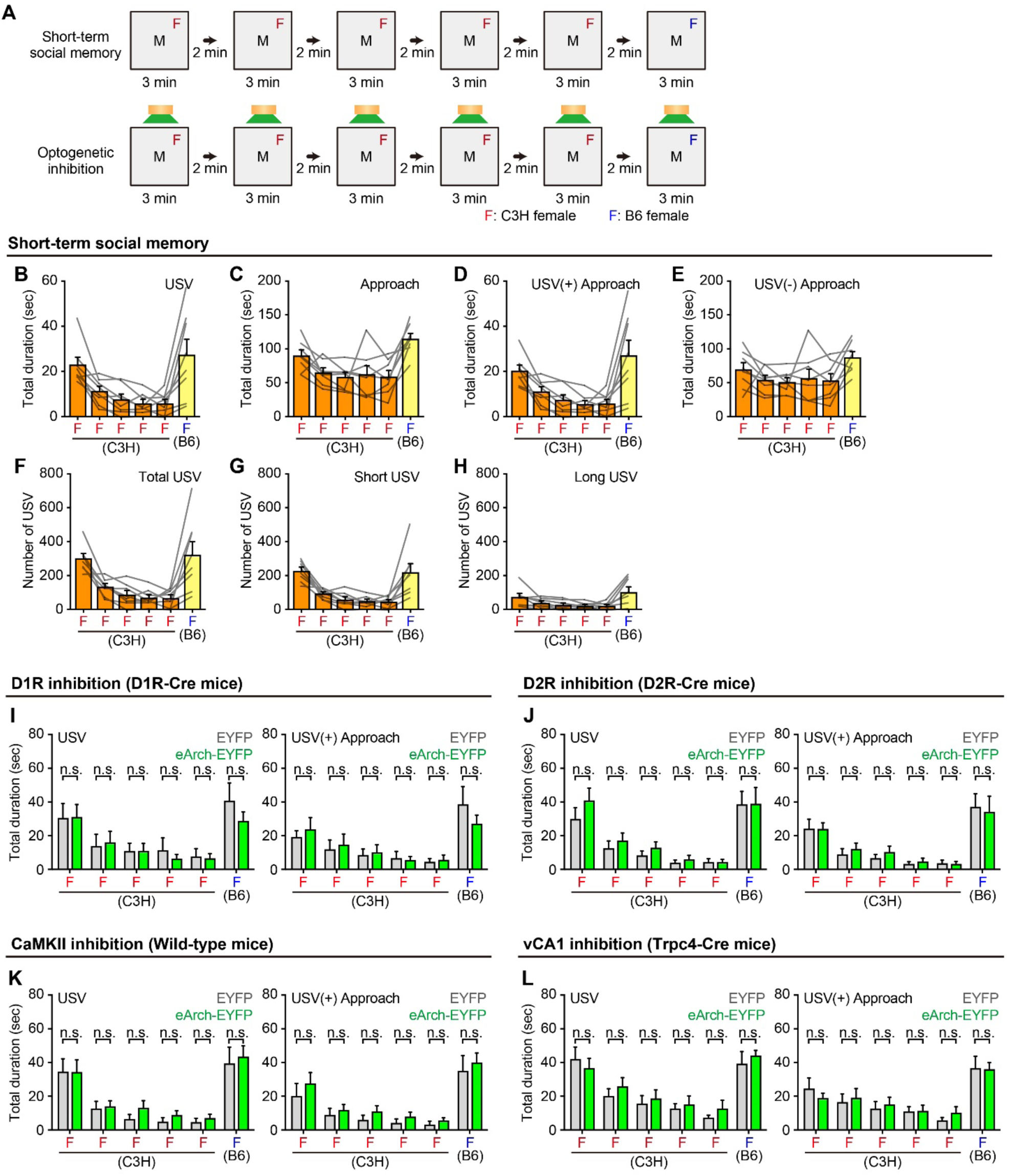
Sexual Behavior Modulation in the Short-term Social Memory Task. **(A)** Schematic view of the short-term social memory task. top: The female mice were introduced to the male mice (black M) for three minutes in two-minute intervals. Interaction with a C3H female (red F) was repeated five times, followed by an interaction with B6 female (blue F). bottom: The short-term social memory task with optogenetic inhibition. **(B, C)** Total duration of USV **(B)** and approaching behaviors **(C)** during the short-term social memory task. n = 10 mice. See source data file for statistical information. **(D, E)** Total duration of approaching behaviors with **(D)** and without **(E)** USVs during the short-term social memory task. See source data file for statistical information. **(F–H)** Total duration of total **(F)**, short **(G)**, and long **(H)** USV during the short-term social memory task. See source data file for statistical information. **(I–L)** Optogenetic inhibition of the NAc-Drd1 **(I)**, NAc-Drd2 **(J)**, NAc-CaMKII **(K)**, and vCA1 **(L)** of male mice during the short-term memory task. Total duration of USVs (left) and approaching behaviors with USVs (right). **(I)** EYFP, n = 8; eArch-EYFP, n = 8 mice. **(J)** EYFP, n = 8; eArch-EYFP, n = 7 mice. **(K)** EYFP, n = 8; eArch-EYFP, n = 9 mice. **(L)** EYFP, n = 7; eArch-EYFP, n = 9 mice. EYFP vs eArch-EYFP, Trial 1-Trial 5: Two-way rANOVA followed by Tukey–Kramer post hoc comparisons, Trial 5-Trial 6: Three-way rANOVA followed by Tukey–Kramer post hoc comparisons.

Furthermore, to investigate the role of D1R-, D2R-, and CaMKII-expressing neurons in the NAc and hippocampal vCA1 neurons in the decay of USV quantity associated with the formation of social memory, we performed optogenetic inhibition of the target neurons during the short-term social memory task, similar to the previous experiments (***Figure 2–Figure 4***). Light stimulation for the inhibition was applied during all social interaction sessions (***Figure 7A***). There were no significant differences across all the experiment groups in terms of the total USV duration and total duration of approaching behavior with the presence of USVs (***Figure 7I–7L***, described in more detail in the source data file), suggesting that the vCA1 and the NAc are not essential for the short-term social memory task.

## Discussion

In this study, we found a reduction in the total number of USVs toward a female conspecific after familiarization with the female. Here, we mainly discuss the following three points: (1) the changes in behavior influenced by social memory about females; (2) the modulation of USV emissions by hippocampal vCA1 neurons responsible for storing social memory; and (3) the dynamic alterations of the *in vivo* physiological features of D1R- and D2R-expressing neurons in the NAc.

### Behavioral changes due to social memory about females

First, in this study, we succeeded in quantifying the social memory of male mice about opposite-sex conspecific females by assessing USV emissions, which is a part of stereotypic sexual behaviors. The number of USVs emitted by a male mouse after familiarization with a female was reduced towards the familiar female, but not towards the novel female. Although our results are inconsistent with the effects of familiarization with females on male-female USVs reported in a previous study (Sasaki et al., 2020), their study used females that had lived together for a week as familiar stimulators; therefore, it is possible that endocrine changes caused by mating or extended co-housing periods with the familiar females may have influenced USV emission. In contrast, as males were familiarized with females without mating for a duration of 2 hours, we were able to measure the effects of social memory alone on USVs. Approaching behavior toward a stimulator, which has previously been used to quantify social memory in male mice (Okuyama et al., 2016), was not affected by forming the memory about the stimulator female in our behavioral paradigm (***Figure 1***), as well as in the mating partner preference test in mice (Cymerblit-Sabba et al., 2020). These findings support the idea that USV emissions are a more suitable indicator for quantifying the memory of female individuals in male mice rather than approaching behavior, which has conventionally been used.

### The modulation of USV emissions by the hippocampal vCA1 neurons that store social memory

Secondly, we found that the reduction in USVs due to social memory formation was blocked by inhibiting the vCA1, as well as D1R- and D2R-expressing neurons in the NAc. Although the reduction of USVs dependent on social memory formation was observed in both the long-term (***Figure 1***) and the short-term (***Figure 7***) social memory tasks, optogenetic inhibition of the vCA1 or NAc affected the results of the long-term social memory task (***Figure 2***), but not those of the short-term task (***Figure 7***). These findings suggest that the functions of the vCA1 and NAc were indispensable for the retrieval of long-term social memory and/or memory based-behavioral changes. We previously reported that hippocampal vCA1 neurons store social memory about same-sex conspecifics and the excitatory neural projection from the vCA1 to the NAc modulate the social interaction after social memory formation (Okuyama et al., 2016). In our novel social memory task, the degree of USV reduction became significantly smaller when the vCA1 was optogenetically inhibited during the retrieval phase after familiarization, probably because of the blurred discrimination between familiar and novel females. Taken together, our study suggests that the vCA1 stores social memory about females as well as males, which is crucial for the social memory based-behavioral changes, such as reduced social interaction with familiar males and reduced USV emissions toward familiar females.

A series of studies has revealed two essential hippocampal subregions that play a role in social memory representation, namely the dorsal CA2 (dCA2) for information processing and its downstream vCA1 for the storage of social memory (Watarai et al., 2021). Among the studies, it was shown that the dCA2 is indispensable for both long-term (Hitti and Siegelbaum, 2014; Meira et al., 2018) and short-term tasks (Hitti and Siegelbaum, 2014). The neural circuit from dCA2 to vCA1 has been reported to form the basis of long-term social memory via sharp-wave ripples (Meira et al., 2018; Rao et al., 2019; Oliva et al., 2020; Tao et al., 2022), suggesting that different circuits from the dCA2 to other regions such as the lateral septum (Lukas et al., 2013; Menon et al., 2022; Rizzi-Wise and Wang, 2021; Leroy et al., 2018) and/or the dorsal CA1 (Chai et al., 2021), that is independent of the vCA1, may function in the short-term social memory task with a few minutes of inter-trial intervals.

### The dynamic alteration of the *in vivo* physiological features in D1R- and D2R-expressing NAc neurons

Our fiber photometry recordings in the NAc revealed that the activities of both D1R- and D2R-expressing neurons were enhanced at the onset of social interaction with stimulator females. Furthermore, the activities of D1R and D2R neuronal populations during the social interaction with familiar or novel females were altered after the familiarization: the activity of D1R-neurons was reduced only in response to familiar females. In contrast, although responses of D2R-neurons were reduced for both familiar and novel females after social memory formation, a significantly stronger reduction was observed toward novel females compared to familiar ones.

In mice and rats, social interaction with females has been shown to enhance dopamine (DA) release in the NAc, as measured by microdialysis (Beny-Shefer et al., 2017; Wenkstern et al., 1993). The release of DA in males decreases with continuous exposure to the same memorized female, corresponding with a decrease in sexual motivation. The release of DA recovers upon exposure to a novel female, leading to the restoration of sexual motivation (i.e., the Coolidge effect) (Fiorino et al., 1997). Recent studies using fiber photometry combined with a genetically encoded fluorescent DA sensor clearly demonstrated that the DA is released in the male NAc at the onset of sexually motivated approach toward females (Dai et al., 2022), which may have resulted in the increased activity of D1R- and D2R-expressing neurons at the onset of social interaction observed in our study. Additionally, Dai et al. also showed that DA release in the NAc was lower during interactions with a same-sex familiar stimulator compared to a same-sex novel stimulator (Dai et al., 2022). Our study demonstrated antagonistic and different dynamics of *in vivo* neuronal activities of D1R- and D2R-expressing neurons in the NAc when interacting with opposite-sex familiar and novel stimulators.

It is well known that D1R- and D2R-expressing neurons in the NAc have different and often antagonistic roles with respect to reward and social behaviors (Soares-Cunha et al., 2016; Cole et al., 2018; Le Merrer et al., 2023). Dopamine-deficient (DA-/-) mice exhibit impairments in sexual behavior, but the administration of a D1R agonist, but not a D2R agonist, to the DA-/- mice leads to an improvement in these impairments (Szczypka et al., 1998). Additionally, D1R-MSN ablation in the NAc results in a decrease in sexual arousal, while D2R-MSN ablation increases sexual arousal (Detraux et al., 2021). Taken together, the D1R-MSNs in the NAc play a stimulatory role in mouse sexual behavior, while the D2R-MSNs play an inhibitory role. In the case of prairie voles, when males formed social memories of their mating partners, males prefer to socially interact with the familiar females over unfamiliar ones and exhibit increased aggression towards unfamiliar females (Aragona et al., 2006). During social memory formation, DA release in the NAc is stimulated by mating, and then the expression of D1R in the NAc neurons was subsequently enhanced (Edwards and Self, 2006; Aragona and Wang, 2009). A series of pharmacological studies has revealed that the administration of a D2R antagonist to the NAc before social memory formation results in the loss of preference behavior for the mating partner after mating (Gingrich et al., 2000; Aragona et al., 2003). Conversely, the administration of a D1R antagonist to the NAc after memory formation leads to an impairment of the decreased aggression towards unfamiliar females (Aragona et al., 2006). Taken together, in voles, D2R signaling is essential for social memory formation and/or pair bonding, whereas D1R signaling is required for the maintenance of formed memories and/or the behavioral change after pair bonding (Aragona and Wang, 2009). Most importantly, the dynamic features of the D1R and D2R neuronal activities observed in our study can be explained by both physiological features related to (1) D1R and D2R antagonistic alteration correlating with sexual motivation and (2) non-specific D2R attenuation, after the formation of social memory about a female (***Figure 8*)**. Regarding the former (1), when encountering the opposite-sex familiar female, the activity of D1R-neurons decreases, while the activity of D2R-neurons is enhanced probably due to the mutual inhibition between D1R-MSNs and D2R-MSNs (Burke et al., 2017). Concerning the latter (2), by forming social memories, the neural activity of D2R-neurons toward both familiar and novel females decreases, probably because of reduced DA release and/or decreased excitability of D2R-MSNs caused by enhanced D2R (Gai) signaling during social memory formation (Bromberg-Martin et al., 2010; Dai et al., 2022).

**Figure 8.**
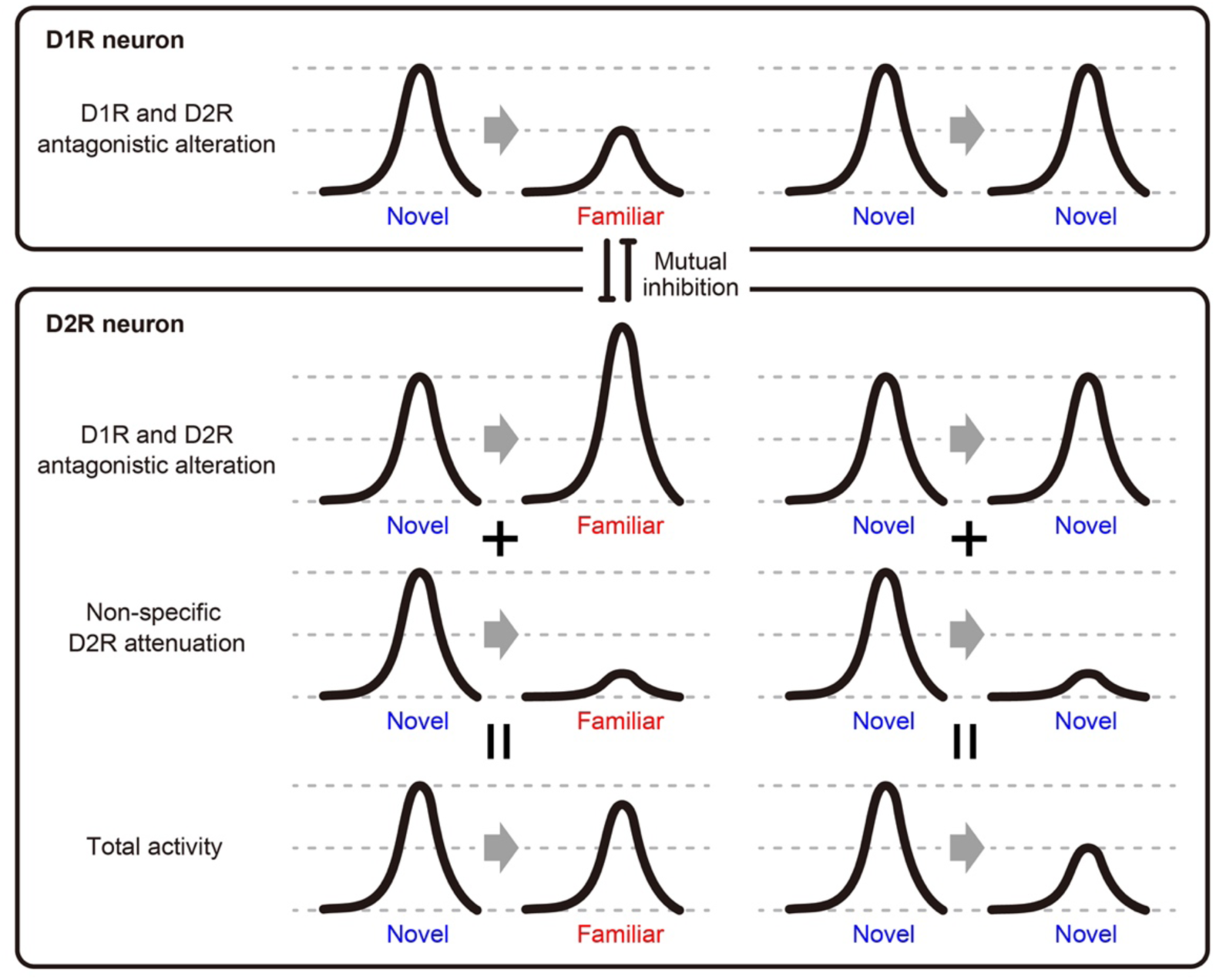
Possible Neural Model.

CaMKII-expressing neurons in the NAc showed enhanced activities to the familiar and the novel females in both the pre- and post-familiarization recordings with no physiological alteration associated with social memory formation (***Figure 6G–6H)***. Furthermore, optogenetic inhibition of the neurons had no effect on social discriminatory behavior (***Figure 4)***, suggesting that D1R- and D2R-expressing neurons, but not CaMKII-expressing neurons in the NAc, were responsive to the behavioral alternation due to the memory about female conspecifics. Considering that optogenetic inhibition was effective only in the long-term task (***Figure 3*** and ***Figure7***), similar to the vCA1 (***Figure 2***), the physiological alteration in D1R- or D2R-expressing neurons, without direct involvement of CaMKII activity, mainly contributes to the social memory-based behavioral modulation in this task. D1R and D2R homodimers signal *through* Gas and Gai cascades to regulate cyclic AMP through the upregulation and downregulation of adenylyl cyclase activity, respectively, whereas the D1R-D2R heterodimer signals through Gaq, phospholipase C, resulting in the enhanced activity of CaMKII through the increased production of IP3 (Rashid et al., 2006; Ng et al., 2010; Osinga et al., 2017). The D1R-D2R heterodimers were found in 20-30% of D1R-expressing neurons in the NAc, with co-expression of CaMKII (Hasbi et al., 2009; Hasbi et al., 2020). Consistent with this evidence, when we visualized D1R, D2R, and CaMKII expression using AAV and transgenic mice lines, we found that co-expression of D1R-CaMKII and D2R-CaMKII occurred in only 20% of D1R- and D2R-expressing neurons, respectively (***Figure 5***). Our study provides insights into the molecular mechanisms underlying social memory based behavioral alternation in the NAc, namely the significant contribution of the D1R and D2R homodimers, but not D1R-D2R heterodimers.

## Material & Method

### Mice

All procedures were performed in accordance with protocols approved by the Institutional Animal Care and Use Committee at the Institute for Quantitative Biosciences, The University of Tokyo. All animals were socially housed in 12 h (7 a.m.–7 p.m.) light/dark cycle conditions with *ad libitum* access to food and water. We used B6 (C57BL/6J, Clea Japan, Inc.), C3H (C3H/HeJJcl, Clea Japan, Inc.), D1R-Cre (B6.FVB(Cg)-Tg(Drd1a-cre)FK150Gsat/Mmucd, 036916-UCD), D2R-Cre (B6.FVB(Cg)-Tg(Drd2-cre)ER44Gsat/Mmucd, 032108-UCD), and Trpc4-Cre mice in the present study. D1R-Cre, D2R-Cre, and Trpc4-Cre mice are characterized by Cre recombinase expression driven by promoters of CNS-specific genes using bacterial artificial chromosome (BAC) constructs. BAC-Cre constructs are inserted at the initiating ATG codon in the first coding exon of the D1R- and D2R-Cre mice and at the translation initiation site in the second coding exon of the Trpc4-Cre mice, respectively (Gong et al., 2007; Okuyama et al., 2016).

### The preparation and equipment for behavioral and ultrasonic vocalization recording

To attenuate ultrasonic noise, we prepared a soundproof box (Danbocchi, KANDA PACKAGE) and placed an acrylic box (38 cm × 38 cm) inside as a test chamber. Under dark conditions, we performed behavioral recording and tracked the body positions of three key points (nose, center, and tail) using an infra-red (IR) sensitive GIG-E camera with two IR illuminators, which was facilitated using Ethovision XT software (Noldus) with an interaction module. Ultrasonic sounds were recorded using an ultrasound microphone (CM16/CMPA, Avisoft-Bioacoustics) and an ultrasound recording interface (UltraSoundGate 116H, Avisoft-Bioacoustics). Using a MATLAB-based algorithm, USVSEG (ver. usvseg09r2) (Tachibana et al., 2020), ultrasonic vocalizations (USVs) were defined as syllables extracted from recorded ultrasonic sounds within the range of 40 kHz - 160 kHz, lasting between 3 msec - 300 msec, and having an intensity greater than 4.5 times standard deviations (SD).

USVs toward females are not always observed in males. However, when male mice engage in interactions with females, sexual experiences during the interaction improves the proportion of individuals who emit USVs toward females (Kanno and Kikusui, 2018). Therefore, all male mice used in the behavioral experiments gained sexual experience through co-housing with a female for at least one night before turning 12 weeks of age. After isolation for at least 2 days, we exposed a B6 female to the male mice and recorded their USVs. Males emitting USVs were used for further behavioral experiments, and not emitting males were excluded from the experiments.

### Long-term social memory assay

We performed the long-term social memory assay in the test chamber, using WT, D1R-Cre, D2R-Cre, and Trpc4-Cre mice. The long-term social memory assay was performed according to previously described methods with some modifications (Okuyama et al., 2016). Briefly, the assay consisted of four phases: Before, Familiarization, Isolation, and After. In the Before and the After phases, we exposed a male test mouse to a C3H or B6 female and recorded his behaviors and USVs for 3 minutes. Next, after a 2-minute interval, we exposed the male to another female for 3 minutes. To avoid bias, the order of presentation was randomized. Familiarization was performed in the test male’s home cage for 2 hours. Considering that USVs are no longer emitted during the post-ejaculatory refractory period (PERP), which lasts for several days in C57BL/6J mice (Valente et al., 2021), the test mouse was familiarized with a C3H female trapped in a pencil holder to prevent any sexual interaction between the test mouse and the female that could lead to ejaculation. Then, the female was removed from the home cage, and the test mouse remained alone for 30 minutes in the Isolation phase. In the long-term social memory assay combined with optogenetic inhibition of neuronal cell bodies using eArch-EYFP, a 561 nm green laser (MGL-N-561-300mW, CNI) was continuously active for 3 minutes during each recording trial and was turned off for the 2 minute intervals. At the beginning of every experiment, the laser output was tested to confirm the delivery of 10 – 12 mW of power to the ends of the optic fiber patch-cords. We divided the 3 minute long trial recordings into 2500 time bins, and the duration of approaching behaviors or USVs was calculated by summing the bins in which each event was detected. The approaching behavior-positive bins were defined as bins in which any of the three body points (nose, center, or tail) of two mice were closer than 5 cm. The number of USVs was defined as the number of syllables detected by USVSEG during the recording period.

### Short-term social memory assay

Similarly, the short-term social memory assay was conducted in the test chamber, using WT, D1R-Cre, D2R-Cre, and Trpc4-Cre mice. This assay consisted of a sequence of six consecutive trials: five trials using a C3H female and one trial using a B6 female. Behaviors and USVs were recorded for a duration of 3 minutes during each trial, with 2-minute intervals between trials. In the short-term social memory assay combined with optogenetic inhibition of neuronal cell bodies with eArch-EYFP, a 561 nm green laser (MGL-N-561-300mW, CNI) was continuously active for 3 minutes during each recording trial and was turned off for the 2 minutes intervals. At the beginning of every experiment, the laser output was tested to confirm the delivery of 10 – 12 mW of power to the ends of the optic fiber patch-cords. We divided the 3 minute long trial recordings into 2500 time bins, and the duration of approaching behaviors or USVs was calculated by summing the bins in which each event was detected. The approaching behavior-positive bins were defined as bins in which any of three body points of two mice were closer than 5 cm. The number of USVs was defined as the number of syllables detected by USVSEG during the recording period.

### Adeno-associated viruses (AAV)

AAV5-CaMKIIa:EYFP and AAV5-CaMKIIa:eArch3.0-EYFP were generated by and acquired from the University of North Carolina (UNC) Vector Core, with a titer of 3.6*10^12 genome copies/ml and 3.4*10^12 genome copies/ml, respectively. AAV5-EF1α:DIO-EYFP and AAV5-EF1α:DIO-eArch3.0-EYFP were generated by and acquired from the University of North Carolina (UNC) Vector Core, with a titer of 3.3*10^12 genome copies/ml and 5.0*10^12 genome copies/ml, respectively. AAV5-hSyn:DIO-H2B-mCherry was generated by and acquired from Vigene Biosciences, with a titer of 2.4*10^13. AAV5-syn:FLEX-jGCaMP8s was generated by and acquired from Addgene, with a titer of 2.8*10^13 genome copies/ml. AAV5-CamKII:Cre-SV40 was generated by and acquired from Addgene, with a titer of 1.5*10^13 genome copies/ml.

### Stereotaxic surgery

The methods used for stereotaxic surgery were described previously (Okuyama et al., 2016). Stereotaxic viral injections and optic fiber implants were all performed in accordance with the Institutional Animal Care and Use Committee at the Institute for Quantitative Biosciences. The three types of mixed anesthetic agents consisted of 75 µg/ml of medetomidine (domitor, Orion Corporation), 400 µg/ml of mitazolam (Sando, SANDOZ), and 400 µg/ml of vetorphale (vetorphale, meiji) and were intraperitoneally administrated at 0.1 ml per 10 g of body weight. Viruses were injected using a glass micropipette attached to a 10 ml Hamilton microsyringe through a microelectrode holder filled with mineral oil. A microsyringe pump and its controller were used to control the speed of the injection. The needle was slowly lowered to the target site and remained for 5 min after the injection. Two screws were placed on the skull surrounding the implant site of each hemisphere to provide additional stability. A layer of adhesive cement was applied followed by dental cement to secure the optical fiber implant. A cap made of the top part of an Eppendorf tube was used to protect the implanted fibers. The incision was closed with sutures. Mice were given 1.5 mg/kg metacam as an analgesic and were kept on a heating pad until fully recovered from the effects of atipamezole (Antisedan, Orion Corporation). Mice were allowed to recover for 2 weeks before all subsequent experiments.

For behavioral experiments, bilateral viral delivery and optic fiber implants were aimed at coordinates relative to Bregma. vCA1 injections and implants were aimed at the following coordinates: AP: - 3.16 mm, ML: ± 3.10 mm, DV: - 4.70 mm and AP: - 3.16 mm, ML: ± 3.10 mm, DV: - 4.55 mm, respectively. NAc virus injections and implants were aimed at the following coordinates: AP: + 1.34 mm, ML: ± 0.60 mm, DV: - 4.30 mm and AP: + 1.34 mm, ML: ± 0.60 mm, DV: - 4.10 mm, respectively. The volumes of AAV5-CaMKIIa:eArch3.0-EYFP and AAV5-CaMKIIa:EYFP used were 150 nl for the NAc of WT mice. The volumes of AAV5-EF1α:DIO-eArch3.0- EYFP and AAV5-EF1α:DIO-EYFP used were 150 nl for the NAc in D1R or D2R-Cre mice or vCA1 in Trpc4-Cre mice. A cocktail of both AAV5-CaMKIIa:EYFP and AAV5-hSyn:DIO-H2B-mCherry was mixed in a 1:1 ratio, and the final volume was 150 nl for the NAc of D1R or D2R-Cre mice. The volume of AAV5-syn:FLEX-jGCaMP8s used was 150 nl for the NAc of D1R or D2R-Cre mice. A cocktail of both AAV5-syn:FLEX-jGCaMP8s and AAV5- CamKII:Cre-SV40 was mixed in a 1:1 ratio, and the final volume was 150 nl for the NAc of WT mice.

### Histology and immunohistochemistry

The methods used for immunohistochemistry were previously described (Okuyama et al., 2016). Briefly, adult mice were transcardially perfused with 4% paraformaldehyde (PFA) in phosphate buffered saline (PBS). Brain sections (50 mm in thickness) were prepared using a vibratome, followed by incubation in 0.3% Triton-X PBS with 5% normal goat serum (NGS) for 1 hour at room temperature (RT). The following primary antibodies were added to the 5% NGS/0.3% Triton-X PBS solution and the tissue sections were incubated overnight at 4°C: chicken anti-GFP (A10262, Invitrogen, 1:1000), mouse anti-tyrosine hydroxylase (TH; MAB318, Millipore, 1:1000), and rabbit anti-red fluorescent protein (RFP; 600-401-379, Rockland, 1:1000). After rinsing with 1× PBS three times for 15 minutes each, tissue sections were incubated with Alexa Fluor-488 or Alexa Fluor-546 conjugated secondary antibodies (Invitrogen, 1:500) for 3 hours at RT. Sections were then washed thrice in 1× PBS for 15 minutes each and mounted using Fluoromount/Plus (Diagnostic BioSystems) on glass slides. All samples were stained by DAPI (1µg/ml) for 15 min. Fluorescence images were capturing by confocal microscopy (FV3000, Olympus) as well as by fluorescence microscopy (BZ-X710, Keyence) using 10X objectives.

The ImageJ plugin StarDist was used to mark regions of interest (ROIs) of RFP positive neurons in confocal images, which were labeled via injection of AAV5-hSyn:DIO-H2B-mCherry into the NAc of D1R-Cre or D2R-Cre mice. To detect CaMKII positive neurons, noise removal (default setting, Despeckle) and background subtraction (30.0 pixels, Rolling Ball Radius) were performed on the channel, and the processed image was converted to a binary image using the default settings of ImageJ. To count the number of mCherry and EYFP double-stained neurons, the ROIs of RFP-positive neurons were applied to the corresponding binarized images, and the ROIs with a mode of 255 were counted as double-stained neurons. Those numbers were normalized by the area size for counting neurons, and the normalized values were used for statical analyses.

### Fiber Photometry

For *in vivo* calcium imaging of CaMKII-, D1R-, or D2R-expressing neurons in the NAc, we injected a CaMKII-GCaMP8s cocktail (150 nl) into the NAc of WT mice and a Cre-dependent GCaMP8s virus (150 nl) into the NAc of D1R-Cre or D2R-Cre mice. For the CaMKII-GCaMP8s cocktail, both AAV5-syn:FLEX-jGCaMP8s and AAV5-CamKII:Cre-SV40 were mixed in a 1:1 ratio. Fiber optic probes (0.5 NA, R-FOC-BL400C-50NA, RWD Life Science) were unilaterally implanted above the NAc (coordinates relative to Bregma: AP: + 1.34 mm, ML: ± 0.60 mm, DV: - 4.15 mm). After 3 weeks of viral expression, mice were handled and habituated by tethering to the fiber optic patch cord (0.37 NA, D207-1479, Doric Lenses) in the test chamber for 10 minutes. The fiber optic patch cord was connected to a fluorescence detector (ilFMC6, Doric Lense) and delivered lights to excite and record the GCaMP signal. Fluorescence excitation was provided by the two connectorized excitation wavelengths (405 nm and 465 nm modulated at 333.786 Hz and 208.616 Hz, respectively), and the emission wavelengths were collected 120 times per second. GCaMP signals were differentiated from the isosbestic signal through the fluorescence detector (Doric neuroscience studio).

Calcium imaging and behavioral results of test mice were recorded for 10−15 minutes and consisted of three female exposures and two intervals lasting for 4 minutes. The female exposure was performed for 30 seconds after the male’s first interaction with the female. The first interaction time points in each exposure were utilized as the behavioral onsets for creating the peristimulus time histogram (PSTH) mentioned below. B6 and C3H females were prepared for the exposure, and C3H females were used for familiarization. Familiarization was performed in the home cage for 2 hours with a C3H female restrained in a pencil holder. The female was then removed, and the test mouse was left in isolation for 30 minutes. Pre- and Post-familiarization recordings were taken before and after the familiarization, and four different recordings (Pre-C3H, Post-C3H, Pre-B6, Post-B6) were taken for each test mouse. The recordings for different strains were not taken on the same day. Note that familiarization for Post-B6 was also performed using C3H females.

The isosbestic signal was scaled to fit the GCaMP signal for the recording session. The fitted isosbestic signal was subtracted from the GCaMP signal and used to normalize the data to determine changes from baseline fluorescence (dF/F). The Z score was obtained by subtracting the mean dF/F from dF/F and dividing it by the standard deviation of dF/F. Furthermore, Z scores were computed within a 5-second window extending from 5 seconds before to 5 seconds after the behavioral onsets for the purpose of calculating the PSTH. The final PSTHs were derived by subtracting the mean value obtained from the 5 sec preceding the behavioral onset from the PSTH. The processing of these signals was performed using GuPPy, a python-based analysis tool for fiber photometry (Sherathiya et al., 2021). The mean of the three final PSTHs in each trial was used for statistical analyses.

Statistical analysis was conducted by calculating the AUC and the peak Z score using the mean of the three final PSTHs in each trial. The AUC was determined from the averaged PSTHs through numerical integration employing the trapezoidal rule, both before and after the behavioral onset.

To calculate the peak Z score, high-amplitude events (local maxima 2 median absolute deviations above the median occurring every 40 msec) were filtered out from the averaged PSTH. The maximum value before and after the onset was determined as the peak Z score.

### Analysis and statistics

All data in the present study were analyzed using MATLAB. To compare behavioral parameters of WT males toward stimulator B6 or C3H female in the long-term social memory assay, the data were analyzed using a two-way repeated-measure analysis of variance (rANOVA) with factors of phase and stimulator type. The behavioral parameters toward stimulator B6 or C3H female in the long-term assay with optogenetic inhibition were analyzed using a three-way rANOVA with factors of phase, stimulator type, and virus type. Furthermore, to compare the EYFP and eArch-EYFP groups, discrimination indices of behavioral frequency or durations were obtained by subtracting the value in the after phase from that in the before phase and dividing it by the sum of both values from the before and after phase. The discrimination indices were analyzed using a two-way rANOVA with factors of stimulator type and virus type. A conservative analysis using Tukey–Kramer correction to account for multiple comparisons in the long-term assay is also shown in the text when appropriate.

In the histological analysis of the neural population in the NAc, the number of double-stained neurons was compared using an unpaired t-test between D1R-Cre and D2R-Cre. The Z scores of dF/F were used for calculating their respective area under the curve (AUC) and peak amplitude in the calcium imaging for long-term memory. Those values were analyzed using a three-way rANOVA (with the factors onset, phase, and stimulator type) in each neural population: D1R, D2R, CaMKII. A conservative analysis using Tukey–Kramer correction to account for multiple comparisons in the calcium imaging is also shown in the text when appropriate. In the short-term social memory assay using WT mice without brain surgery, the behavioral parameters toward C3H females during the five consecutive trials were analyzed by rANOVA with the factor of trial. Further, we compared the behavioral parameters between C3H females in the fifth trial and B6 females in the sixth trial using a paired t-test. To compare the EYFP and eArch-EYFP groups in the short-term social memory assay with optogenetic inhibition, the behavioral parameters were analyzed by a two-way rANOVA with factors of trial and virus type. A conservative analysis using Tukey–Kramer correction to account for multiple comparisons in the short-term assay is also shown in the text where appropriate.

## Acknowledgement

We thank I. Yoshimura, M. Sako, A. Watarai, and M.T. Tang for technical assistance, K. Kanno, R. Tachibana, and M. Kato for advising USVs recording setup and analysis methods, and all members of the Okuyama laboratory for discussion and support. This work was supported by JST (Grant Numbers JPMJFR2143 and JPMJCR23B1 (to T.O.)), JSPS KAKENHI (Grant Numbers JP20J01468 (to A.W.), JP18H02544, JP20K21459, JP21H02593, and JP21H05140 (to T.O.)), AMED (Grant Number JP21wm0525018 (to T.O.)), the Naito Foundation (to T.O.), and SECOM Science and Technology Foundation (to T.O.).

## Author contribution

Conceptualization, A.W., K.I., and T.O.; Methodology, A.W., K.I., and T.O.; Software, A.W. and K.I.; Validation; Formal analysis, A.W. and K.I.; Investigation, A.W. and K.I.; Resources, A.W. and K.I.; Data curation, A.W., K.I., and T.O.; Writing – original draft preparation, A.W., K.I., and T.O.; Writing – review & editing, A.W., K.I., and T.O.; Supervision, T.O.

---

**Figure 2–figure supplement 1.**
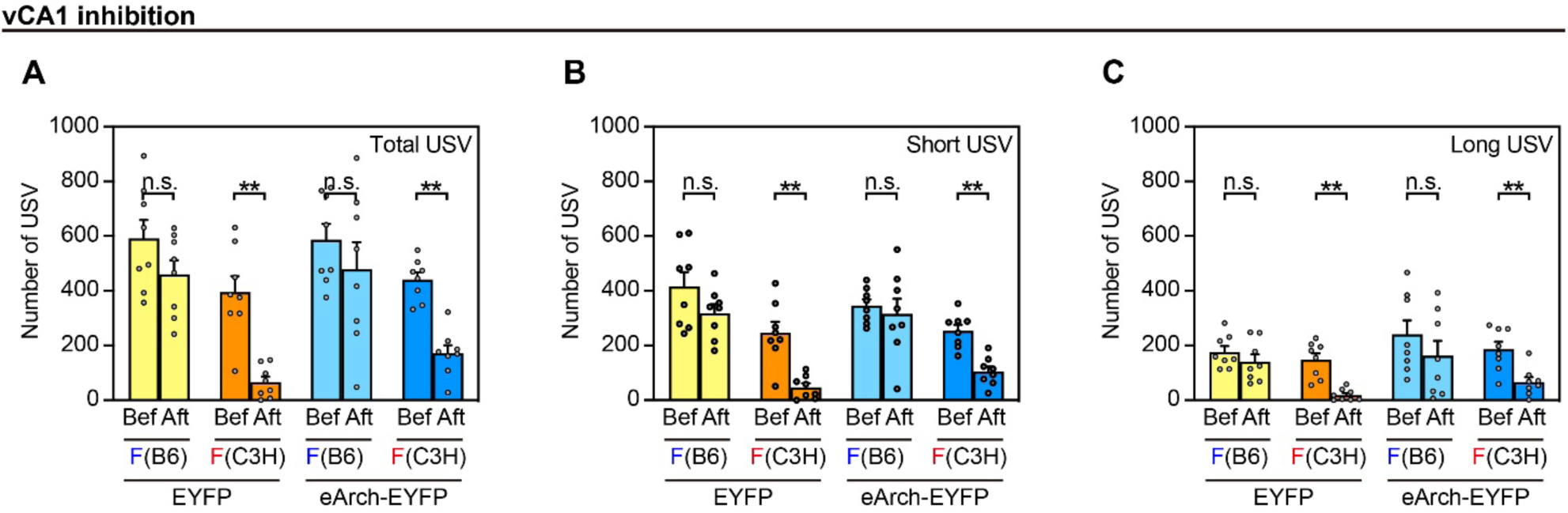
The Number of USVs Emitted During the Long-term Memory Task with Optogenetic Inhibition of vCA1 neurons. **(A–C)** The number of total **(A)**, short **(B)**, and long **(C)** USVs emitted by the male toward B6 and C3H females with and without inhibition of the vCA1. EYFP, n = 8; eArch-EYFP, n = 8. **p<0.01. For all panels: Three-way rANOVA followed by Tukey–Kramer post hoc comparison.

**Figure 3–figure supplement 1.**
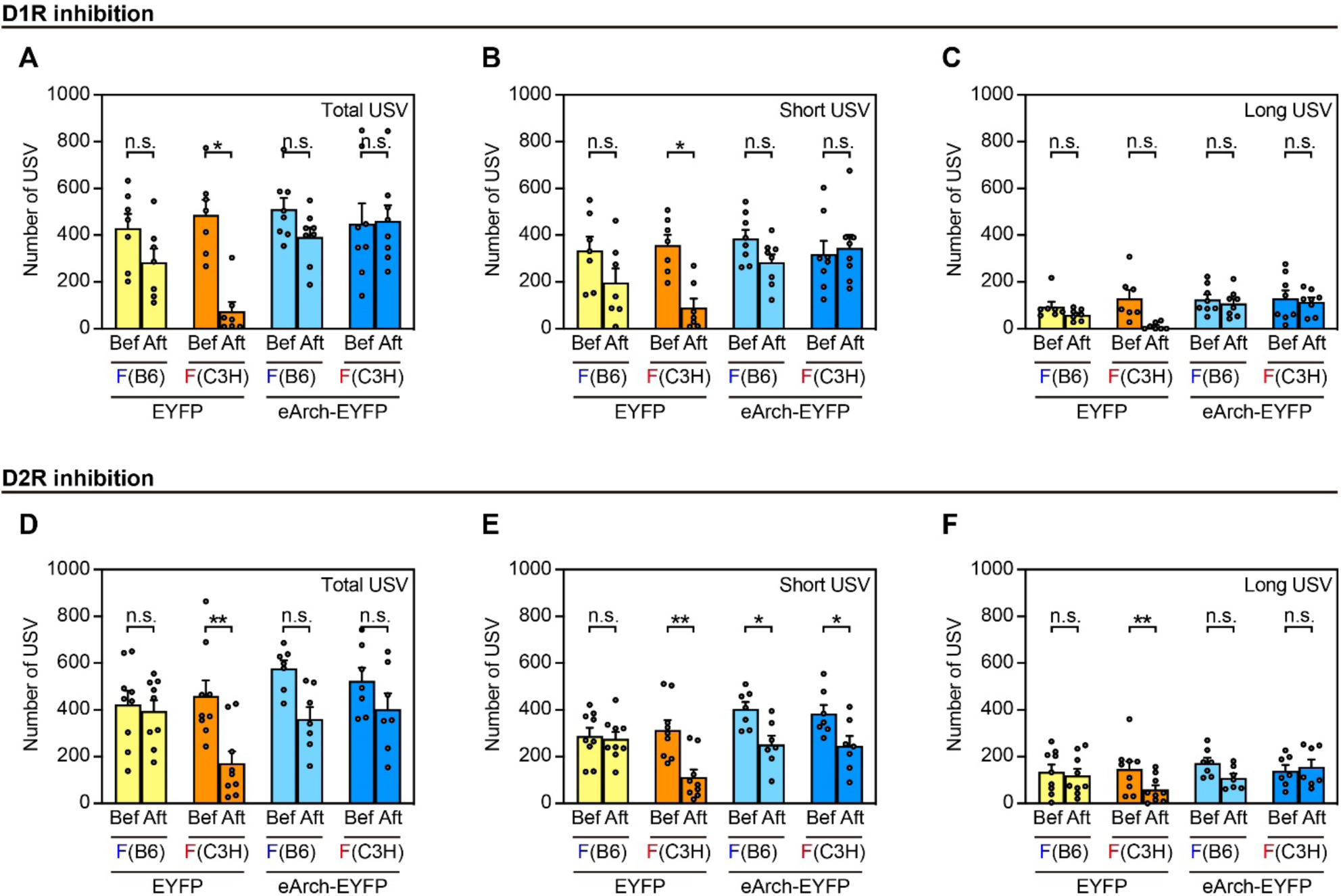
The Number of USVs Emitted During the Long-term Memory Task with Optogenetic Inhibition of D1R- and D2R-expressing NAc neurons. **(A–C)** The number of total **(A)**, short **(B)**, and long **(C)** USVs emitted by the male toward B6 and C3H females with and without inhibition of the NAc-D1R. EYFP, n = 7; eArch-EYFP, n = 8. *p<0.05. **(D–F)** The number of total **(D)**, short **(E)**, and long **(F)** USVs emitted by the male toward B6 and C3H females with and without inhibition of the NAc-D2R. EYFP, n = 9; eArch-EYFP, n = 7. *p<0.05; **p<0.01.

**Figure 4–figure supplement 1.**
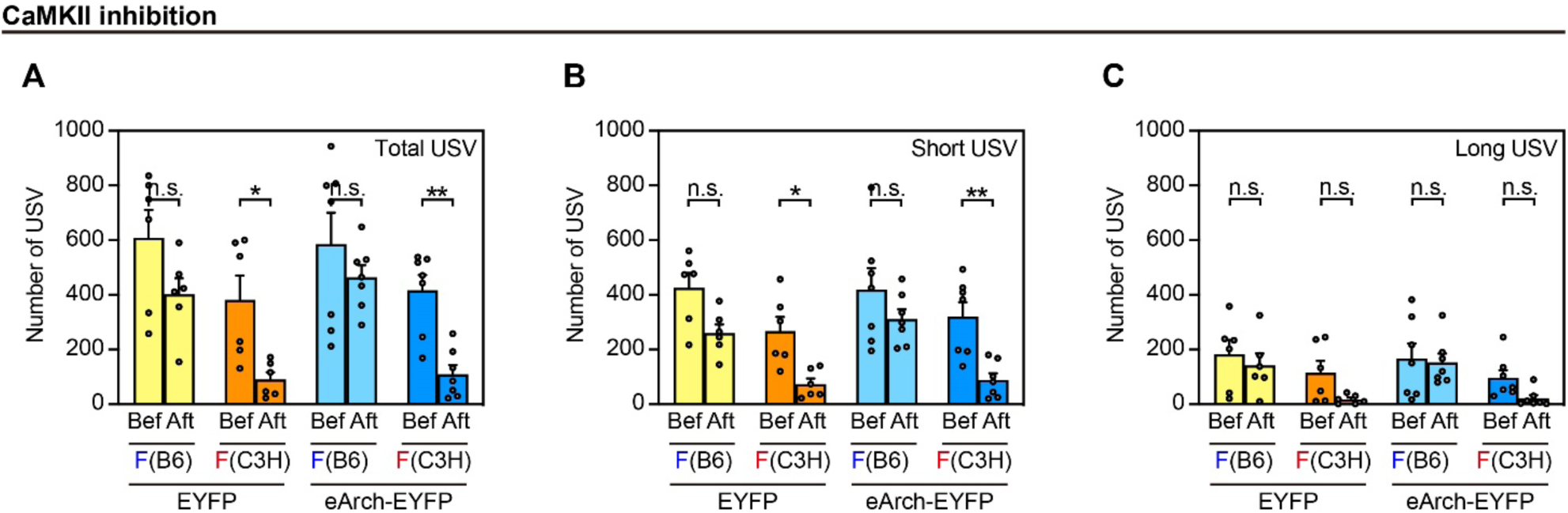
The Number of USVs Emitted During the Long-term Memory Task with Optogenetic Inhibition of CaMKII-expressing NAc neurons. **(A–C)** The number of total **(A)**, short **(B)**, and long **(C)** USVs emitted by the male toward B6 and C3H females with and without inhibition of NAc-CaMKII. EYFP, n = 6; eArch-EYFP, n = 7. *p<0.05; **p<0.01.

